# TYK2 as a novel therapeutic target in Alzheimer’s Disease with TDP-43 inclusions

**DOI:** 10.1101/2024.06.04.595773

**Authors:** Laura E. König, Steve Rodriguez, Clemens Hug, Shayda Daneshvari, Alexander Chung, Gary A. Bradshaw, Asli Sahin, George Zhou, Robyn J. Eisert, Federica Piccioni, Sudeshna Das, Marian Kalocsay, Artem Sokolov, Peter Sorger, David E. Root, Mark W. Albers

## Abstract

Neuroinflammation is a pathological feature of many neurodegenerative diseases, including Alzheimer’s disease (AD)^1,2^ and amyotrophic lateral sclerosis (ALS)^3^, raising the possibility of common therapeutic targets. We previously established that cytoplasmic double-stranded RNA (cdsRNA) is spatially coincident with cytoplasmic pTDP-43 inclusions in neurons of patients with C9ORF72-mediated ALS^4^. CdsRNA triggers a type-I interferon (IFN-I)-based innate immune response in human neural cells, resulting in their death ^4^. Here, we report that cdsRNA is also spatially coincident with pTDP-43 cytoplasmic inclusions in brain cells of patients with AD pathology and that type-I interferon response genes are significantly upregulated in brain regions affected by AD. We updated our machine-learning pipeline DRIAD-SP (Drug Repurposing In Alzheimer’s Disease with Systems Pharmacology) to incorporate cryptic exon (CE) detection as a proxy of pTDP-43 inclusions and demonstrated that the FDA-approved JAK inhibitors baricitinib and ruxolitinib that block interferon signaling show a protective signal only in cortical brain regions expressing multiple CEs. Furthermore, the JAK family member TYK2 was a top hit in a CRISPR screen of cdsRNA-mediated death in differentiated human neural cells. The selective TYK2 inhibitor deucravacitinib, an FDA-approved drug for psoriasis, rescued toxicity elicited by cdsRNA. Finally, we identified CCL2, CXCL10, and IL-6 as candidate predictive biomarkers for cdsRNA-related neurodegenerative diseases. Together, we find parallel neuroinflammatory mechanisms between TDP-43 associated-AD and ALS and nominate TYK2 as a possible disease-modifying target of these incurable neurodegenerative diseases.

## INTRODUCTION

Efficacious therapies for cancer patients based on principles of precision medicine have impacted millions of patients suffering from this heterogenous disease^5–7^. Mounting evidence indicates that clinical AD, like cancer, is a collection of multifaceted neurodegenerative diseases^8–10^. With the number of AD patients increasing in our aging society^11^, it is likely that personalized treatments for specific subsets of AD will be important to address this huge unmet need for effective therapy.

Pathologic features that stratify AD – a disease defined by amyloid beta (Aβ) plaques and neurofibrillary tangles (NFTs) of hyperphosphorylated tau protein – are proteinopathies and include the formation of cytoplasmic inclusions of phosphorylated TAR DNA-binding protein 43 (pTDP-43)^12^. TDP-43-pathology is observed in up to 57% of AD patients^13^ and is the most prominent proteinopathy after Aβ and tau pathology^14^. TDP-43 is a nuclear protein involved in RNA processing^15^. Ectopic cytoplasmic inclusions of this protein are thought to deprive the nucleus of TDP-43 function and dysregulate RNA processing and expression^16–20^. For instance, loss of function of TDP-43 triggers derepression of immunostimulatory genomic double-stranded RNA (dsRNA) that can activate pattern recognition receptors (PRRs)^21,22^, which results in increased cell autonomous and non-cell autonomous inflammatory processes^23^. Previously, we demonstrated that cytoplasmic pTDP-43 inclusions are spatially coincident with cytoplasmic dsRNA (cdsRNA) in ALS and frontotemporal dementia (FTD) patients with an intronic expansion of hexanucleotide repeats in the *C9ORF72* gene^4^. CdsRNA triggered an interferon-mediated toxic innate immune response in a sequence-independent fashion and led to neuronal cell death in in-vitro and mouse models. Elucidating the role of cdsRNA in the setting of neuronal TDP-43-pathology in other neurodegenerative diseases could offer a paradigm to target common neuroinflammatory mechanisms using principles of precision medicine.

Here, we show that cdsRNA and cytoplasmic pTDP-43 inclusions are spatially coincident in brains of AD patients and demonstrate the induction of interferon-signaling in AD patients’ brains. Our previously published machine learning pipeline for drug repurposing in AD (DRIAD-SP) has identified baricitinib and ruxolitinib, both FDA-approved JAK kinase inhibitors, among the top repurposing candidates^24^. Here, we demonstrate that the dominant protective signal for those drugs is exhibited only for a subset of AD patients with elevated CE expression, which is a proxy for TDP-43 pathology. To evaluate the mechanism of action of these compounds, we conducted a CRISPR screen and observed that inhibition of TYK2, a JAK kinase family member, robustly rescues cdsRNA-induced toxicity in a vigorous manner. In addition to baricitinib and ruxolitinib we also found that deucravacitinib – a selective TYK2 inhibitor recently approved for moderate-to-severe plaque psoriasis^25^ – exhibited a neuroprotective effect at a substantially lower IC_50_ compared to the other two drugs. Finally, we analyzed potential inflammatory biomarkers for dsRNA-mediated neuropathology and found that CCL2, CXCL10, and IL6 are candidate biomarkers for validation in clinical trials. Our work, together with an independent work by the Zoghbi laboratory demonstrating that TYK2 phosphorylates tau^26^, supports TYK2 as a candidate disease-modifying drug target.

## RESULTS

### Spatial coincidence of cdsRNA and pTDP-43

To assess the presence of cdsRNA in neurons with cytoplasmic pTDP-43 inclusions in patients with AD, we stained formalin-fixed paraffin-embedded (FFPE) amygdala sections of human post-mortem brains of healthy individuals (n = 9) and of patients who met the pathologic diagnosis of AD (n = 10) (Fig. 1a, Extended Data Table 1). We detected cdsRNA in all AD patients with severe pTDP-43 pathology by immunostaining. In contrast, both pTDP-43 inclusions and cdsRNA were absent in healthy controls (Extended Data Fig. 1a,b). In AD patients with mild pTDP-43 pathology, we detected cdsRNA in 33% of cases. Next, we evaluated the spatial coincidence of cdsRNA and pTDP-43 by performing cyclic immunofluorescence (CyCIF)^27^ which enabled multiplexed staining of pTDP-43 inclusions, cdsRNA, NFTs, Aβ plaques, and nuclei on the same section (Fig. 1b, lower panel of healthy control and AD, respectively). We compared the acquired images to luxol fast blue-hematoxylin and eosin (LHE) stainings as well as immunohistochemical stainings of cdsRNA, pTDP-43, tau, and Aβ (Fig. 1b, upper panel of healthy control and AD, respectively). While NFTs and Aβ plaques were detected in the tissue, neither pathology overlapped spatially with cytoplasmic pTDP-43 inclusions nor cdsRNA (Fig. 1b). However, pTDP-43 and cdsRNA were spatially coincident (enriched two-fold in AD cases; Extended Data Fig. 1c), paralleling our previous observations in ALS brains^4^. To establish a direct link between cdsRNA and activation of PRRs in AD patient’s brains, we immunostained amygdala sections for phosphorylated protein kinase R (PKR), a kinase that autophosphorylates when bound to dsRNA^28^ (Extended Data Fig. 2). We observed that PKR is only activated in pTDP-43/cdsRNA-positive regions of AD brains.

**Fig. 1:**
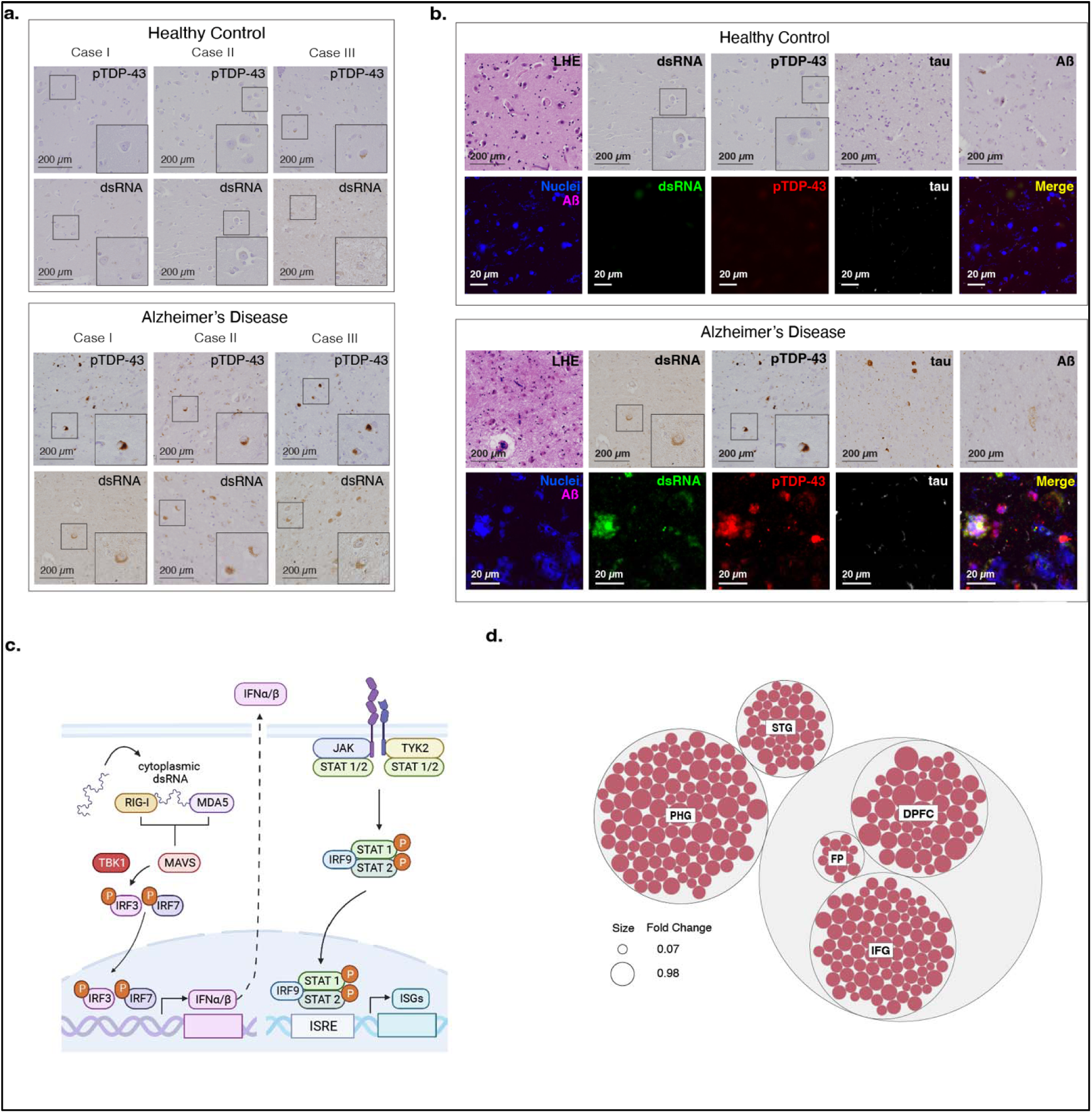
CdsRNA induces IFN-I signaling and is spatially coincident with pTDP-43 inclusions in AD. **a**, Immunohistochemistry (IHC) for cdsRNA and pTDP-43 inclusions in human postmortem brain sections of the amygdala. Top panel: healthy control cases, bottom panel: AD patient cases. Zoom-ins with 2x magnification. **b,** LHE, IHC and cyclic immunofluorescence (CyCIF) of human postmortem brain sections of the amygdala. IHC for cdsRNA, pTDP-43, tau, and Aß on different brain sections of the same healthy control (top panel, healthy control case II)/AD patient (bottom panel, AD case I). CyCIF panels show nuclei (Sytox Blue), Aß, dsRNA, pTDP-43, and tau as well as a merge of all channels. CyCIF was done on a single brain section of the same healthy control as the IHC images, but of a different AD patient. **c,** Pathway schematic of innate immune response to cdsRNA in AD leading to the expression of ISGs. **d,** Bubble chart of upregulated ISGs within relevant brain regions in AD patients. Each red bubble represents a gene. Bubble size is proportional to fold change of upregulation. Chart is based on a differential gene analysis of RNA-sequencing data derived from the ROSMAP^29^ and MSBB^30^ databases comparing AD patients and healthy controls. Individual gene names are shown in Extended Data Fig. 3. DFPC: dorsolateral prefrontal cortex; FP: frontal pole; IFG: inferior frontal gyrus; PHG: parahippocampal gyrus; STG: superior temporal gyrus. Patient information of human postmortem brain sections that are depicted in this figure can be found in Extended Data Table 1.

If cdsRNA acts as a damage-associated molecular pattern (DAMP) in neurons, then we anticipate it would trigger an IFN-I response through the activation of the PRRs RIG-I and MDA5 (Fig. 1c). We conducted differential gene expression analysis of RNA-sequencing data from five relevant brain regions available from the Religious Order Study/Memory and Aging Project (ROSMAP)^29^ and Mount Sinai Brain Bank (MSBB)^30^ AMP-AD databases to test our hypothesis of elevated interferon signaling. We found 139 unique interferon-stimulated genes (ISGs) that were significantly upregulated and 51 ISGs that were significantly downregulated (adjusted *P*-value < 0.05) in AD patients compared to healthy controls. The parahippocampal gyrus (PHG), a brain region that shows atrophy early on in disease progression^31,32^, had the greatest number of upregulated ISGs, followed by the dorsolateral prefrontal cortex (DPFC) and the inferior frontal gyrus (IFG). In addition, we observed upregulation of ISGs, to a lesser extent, in the superior temporal gyrus (STG) and frontal pole (FP; Fig. 1d, Extended Data Fig. 3, Supplementary Table 1). Cytoplasmic pTDP-43 inclusions have been reported in these brain regions^33^, which supports our hypothesis of cdsRNA-mediated IFN-I induction in these AD-associated brain regions.

### DRIAD-SP predicts efficacy of blocking interferon signaling

We previously published a machine learning framework for Drug Repurposing In Alzheimer’s Disease with Systems Pharmacology (DRIAD-SP)^24^ that links the prediction of disease stage (early-vs. late-stage AD based on the Braak AD staging system^34^) to drug-induced molecular signatures in human neural cells. Through this pipeline, we previously identified baricitinib, ruxolitinib, and tofacitinib, which are FDA-approved JAK inhibitors that inhibit interferon signaling, as drug candidates for repurposing in AD^24^ (Extended Data Fig. 4).

Here, we extend the framework along two directions. First, we improve DRIAD-SP’s predictive power by training and evaluating predictors of disease stage using ordinal ridge regression (Fig. 2a) instead of binary classification, as was done previously. This approach allows for training on the full scale of stage I to stage VI along the Braak spectrum^34^, without the need to stratify patients into “early” and “late” categories. Secondly, to tailor DRIAD-SP to AD patients with pTDP-43 pathology, we used the expression of CEs as a proxy for the presence of pTDP-43 inclusions (Fig. 2b). It has been shown that the pre-mRNAs of stathmin-2 (STMN2)^19,20^ and UNC13A^17,18^ have binding sites for TDP-43 and that TDP-43, when physiologically present in the nucleus, represses CE inclusion in the canonical mRNA of both STMN2 and UNC13A during splicing. pTDP-43 pathology is associated with nuclear hypofunction of TDP-43 which leads to CEs being included in the processed mRNA resulting in a short variant of STMN2 mRNA^19,20^ (STMN2 short) due to a premature stop codon, and two versions of CE-inclusive UNC13A mRNA, which we refer to as UNC13A-CE1 and UNC13A-CE2^17,18^. We posited the presence of these CEs in AD brains as a proxy for pTDP-43 pathology. As about 97% of ALS patients exhibit pTDP-43 inclusions^35^, we tested our approach by checking for the presence of CEs in RNA-sequencing data^36^ from FACS-sorted single neuronal nuclei from the neocortex of deceased ALS patients. We confirmed that nuclei that do not contain TDP-43 have higher reads of the CEs STMN2 short (log2 fold change (LFC) = 3.55; *P* < 0.0001), UNC13A-CE1 (LFC = 4.20; *P* = 0.0002) and UNC13A-CE2 (LFC = 4.05; *P* = 0.0006) relative to nuclei that contain TDP-43 (Extended Data Fig. 5). By contrast, the number of reads of canonical STMN2 were reduced to a smaller degree (LFC = -1.76; *P* = 0.0017) and not reduced for UNC13A (LFC = 0.278; *P* = 0.11) mRNAs.

**Fig. 2:**
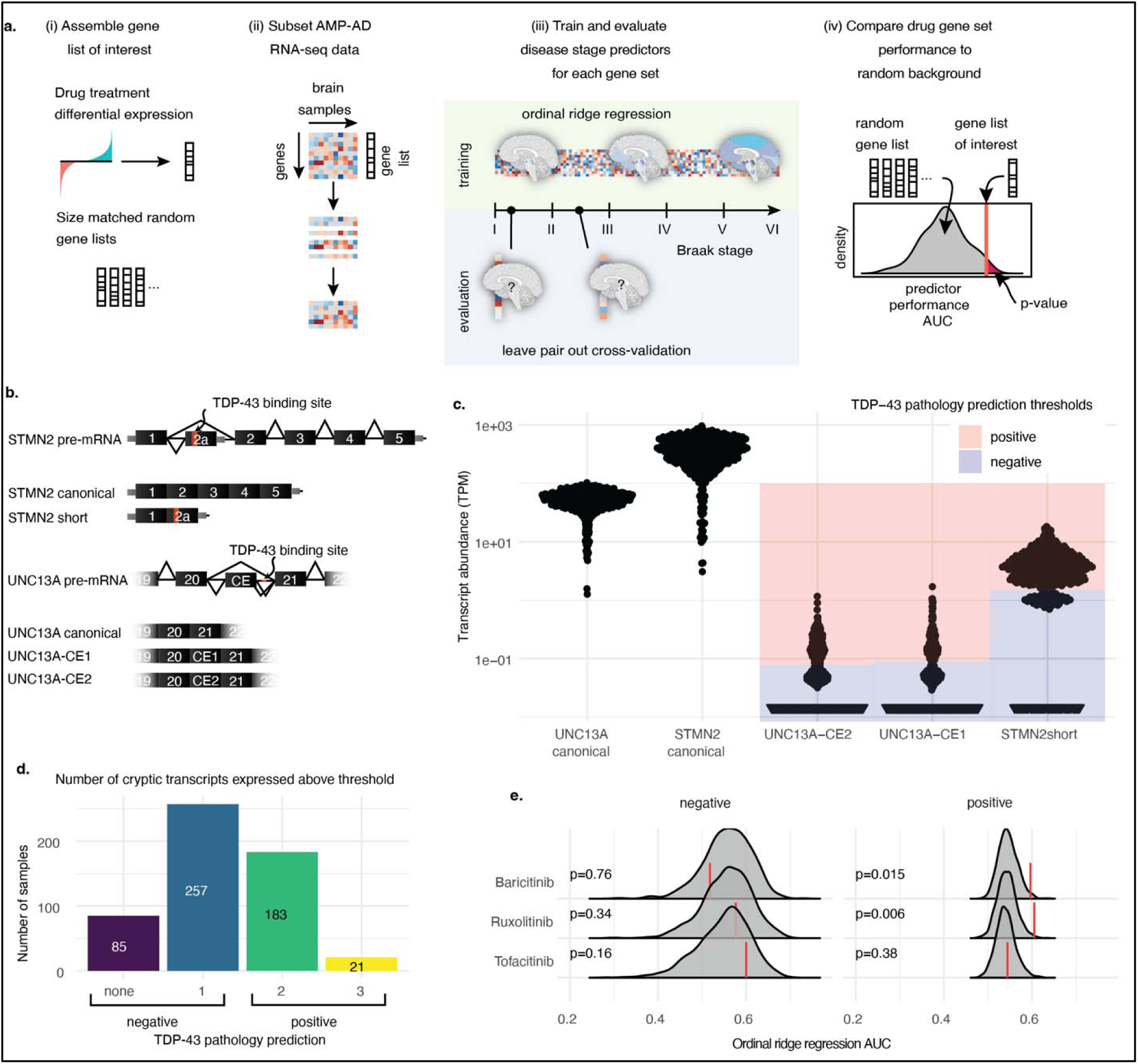
Prediction of drug efficacy in patients stratified by TDP-43 pathology. **a,** Schematic representation of the DRIAD-SP drug efficacy prediction pipeline. From left to right (i) gene sets are assembled from differential gene expression analysis of cell lines treated with the drugs of interest against DMSO controls. For comparison, random gene sets matching the sizes of the drug gene sets are generated. (ii) RNA-sequencing data from AD brain samples is subset to only contain genes that are present in the gene set that is being tested. (iii) The RNA-sequencing subset is used to fit ordinal ridge regression models predicting Braak disease stage of the patient cohorts. Leave pair out cross-validation is used to evaluate model performance (areas under the curve; AUC). (iv) Drug efficacy is predicted by comparing the performance of the model using the drug gene sets to the size-matched random gene sets. **b,** Schematic representation of TDP-43 pathology associated splice variants that include CEs in STMN2 and UNC13A. **c,** Expression of TDP-43 associated splice variants in the PCC brain tissues of the ROSMAP^29^ patient cohort. Blue shaded regions are below, and red regions are above the chosen threshold for expression to be considered positive. **d,** Per-patient-quantification of the number of CE transcripts that are expressed above the chosen thresholds. Patients are predicted to be TDP-43 pathology negative if they express one or fewer of the CE transcripts, and positive if they express two or more. **e,** Performance of DRIAD-SP models for three selected drugs. Drug efficacy was assessed separately in patient populations according to their predicted TDP-43 pathology. Red lines indicate the DRIAD-SP model performance trained on the drug gene sets, whereas the gray shaded regions correspond to the distribution of model performances based on random gene sets.

With this validation, we stratified patients’ RNA-sequencing data derived from ROSMAP and MSBB AMP-AD databases based on the presence of CEs. First, we set the threshold for the presence or absence of each individual CE above the lowest non-zero peak (essentially > 1 CE transcript detected) in the abundance histograms (transcripts per million TPM>1.3 STMN2 short; TPM>0.09 for UNC13A-CE1; TPM>0.08 UNC13-CE2; Fig. 2c). We classified patients with two or more CE transcripts as TDP-43 positive (Fig. 2d). In the MSBB data, the counts for UNC13A-CE1 and UNC13A-CE2 were much lower relative to the ROSMAP data. We reasoned that the single-end sequencing protocol to generate the MSBB RNA-sequencing data was less sensitive for CE detection, relative to the paired-end reads in the ROSMAP data (Extended Data Fig. 6). We restricted our analysis to ROSMAP sequencing data from the posterior cingulate cortex (PCC), a brain region impaired early on in disease progression and therefore highly relevant for our study^37^. Of 546 total patients, no CEs were detectable in 85 patients, and one type of CE was detected in 257 patients; 183 patients had two types of CEs, and 21 patients had all three CEs present (Fig. 2d). Hence, 37% of cases were classified “TDP-43-positive”-AD patients which is a conservative prediction considering that the clinical incidence of TDP-43-pathology in AD patients can reach 57%^13^. An ordinal regression model to see whether the predicted TDP-43 status changes the expected value of the Braak stage resulted in a negative estimate (estimate = -0.17; standard error = 0.16: z value = -1.1), indicating that the CE-positive cases are on average slightly earlier in their disease progression. However, this effect was not significant (*P* = 0.27) meaning that the proportion of patients at different Braak stages was not significantly different between the “TDP-43-positive” and “TDP-43-negative” cases (Extended Data Fig. 7a).

We assessed the predicted efficacy of baricitinib, ruxolitinib and tofacitinib in AD patients with 2 or more CEs (as a proxy of TDP-43 dysfunction) relative to AD patients with one or no CEs (Fig. 2e, further replicates in Extended Data Fig. 7b). Both baricitinib and ruxolitinib, but not tofacitinib, were flagged as significant hits when predictors were trained on RNA-sequencing data from AD patients with 2 or more CEs. On the other hand, none of the three drugs (baricitinib *P* = 0.76, ruxolitinib *P* = 0.34, tofacitinib *P* = 0.16) were significant hits in no / low CE-expressing AD patients. These findings support our hypothesis that there are IFN-I-mediated neuroinflammatory processes occurring in AD patients’ brains with TDP-43 pathology.

### CRISPR screen and validation of TYK2

We replicated cdsRNA-mediated inflammation and death by transfecting differentiated ReN VM cells with poly(I:C), a dsRNA mimetic (dsRNAmi) consisting of polyinosinic-polycytidylic acid. These differentiated human neural cells contain markers for neurons, astrocytes, and oligodendrocytes, but not microglia^38^. In contrast to our previous work (“One Pot” Differentiation, Extended Data Fig. 8a)^4^, we found that differentiating neural cells in a separate dish and then transferring differentiated neurons into the assay plate reduced variability between technical replicates (“Separate Pot” Differentiation; Extended Data Fig. 8b). Both baricitinib and ruxolitinib rescued human neural cell death in this new workflow with greater reproducibility and lower variance (Extended Data Fig. 9a,b) with nearly complete rescue at 1 µM and >100% at 10 µM – presumably due to preventing toxicity from the transfection reagent lipofectamine 2000 present in the control. Both drugs significantly inhibited the phosphorylation of STAT1 and the downstream transcription of ISGs (Extended Data Fig. 9c,d).

Since the EC_50_ of baricitinib (EC_50_ = 35.7 nM) and ruxolitinib (EC_50_ = 47.7 nM) are greater than their affinities to their target proteins in our assay (baricitinib: IC_50_(JAK1) = 5.9 nM, IC_50_(JAK2) = 5.7 nM^39^; ruxolitinib: IC_50_(JAK1) = 3.3 nM, IC_50_(JAK2) = 2.8 nM^40^), we hypothesized that the inhibition of other kinases may be responsible for their neuroprotective effect. To test this hypothesis, we conducted a genome-wide CRISPR screen to rescue cdsRNA-induced toxicity in differentiated human neural cells^4,41^. We used the genome-wide Brunello library which consists of four single guide RNAs (sgRNAs) per human gene and 1,000 control sgRNAs^42^ (Fig. 3a) expressed in barcoded lentiviral particles to transduce ReN VM cells in their neural progenitor state. After a week-long differentiation, one replicate was harvested to identify genes that were essential for ReN cell differentiation and were not present at the initiation of either poly(I:C) or lipofectamine. Eight different genes were significantly underrepresented (*DGAT2, SIGLEC11, GSTA2, ABI1, FBXL8, AHCY, C7orf55-LUC7L2, ARHGEF12*; Supplementary Table 2). After 48 hours of treatment with poly(I:C) or mock lipofectamine transfection, the cells were harvested, genomic DNA was extracted, and the sgRNA sequences within the gDNA were PCR-amplified and sequenced to identify which sgRNA barcodes are enriched. Enriched sgRNAs correspond to candidate target genes whose loss-of-function enhances survival.

**Fig. 3:**
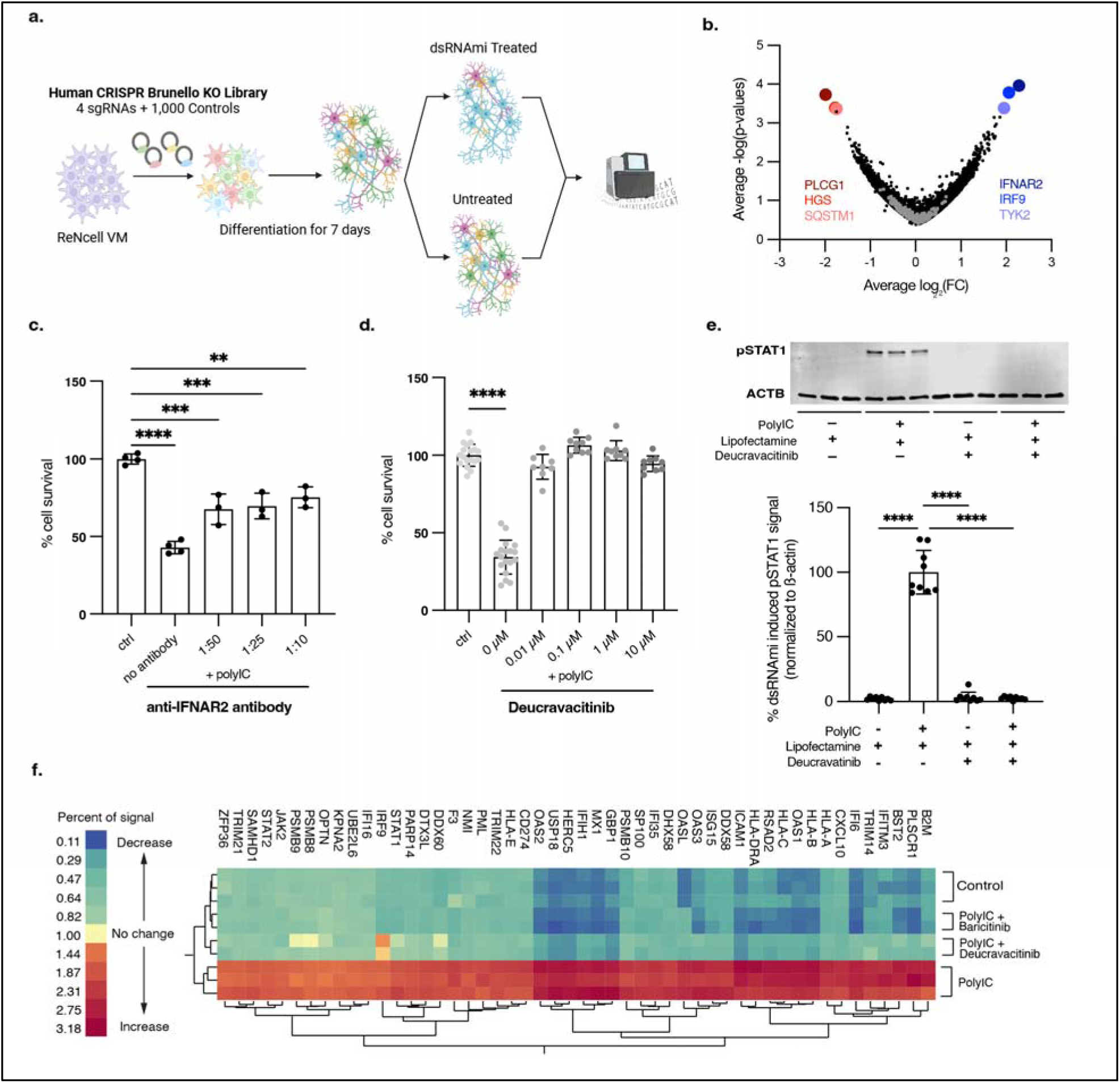
CRISPR screen and subsequent validation identifies TYK2 as a therapeutic target. **a**, Schematic of the CRISPR screen conducted in ReN VM cells using the Brunello library^42^ to identify potential therapeutic targets for rescuing the toxic immune response triggered by cdsRNA. **b,** Volcano plot showing the LFC of genes targeted in the CRISPR screen against their -log(*P*-value) and identifying *IFNAR2* (average LFC = 2.23825, *P* = 0.0001), *IRF9* (average LFC = 2.0625, *P* = 0.0002) and *TYK2* (average LFC = 1.95, *P* = 0.0004) as the most prominent and *PLCG1* (average LFC = -1.9875, *P* = 0.0002), *HGS* (average LFC = -1.775, *P* = 0.0004) and *SQSTM1* (average LFC = -1.7475, *P* = 0.0005) as the least abundant knockouts. Gray dots represent negative control sgRNAs. **c,** Quantification of cell survival of ReN VM cell-derived neurons treated with different concentrations (no antibody *P* < 0.0001, n=4; 1:50 *P* = 0.0001, n = 3; 1:25 *P* = 0.0002, n = 3; 1:10 *P* = 0.0011, n = 3) of anti-IFNAR2 antibody after transfection with poly(I:C) or lipofectamine as vehicle control (ctrl, n = 4). **d,** Quantification of cell survival of ReN VM cell-derived neurons pre-treated with deucravacitinib, a selective TYK2 inhibitor, at different concentrations (0 µM *P* < 0.0001, n = 18; 0.01 µM *P* = 0.1273, n = 9; 0.1 µM *P* = 0.1920, n = 9; 1 µM *P* = 0.8455, n = 9; 10 µM *P* = 0.3562, n = 9) and afterwards transfected with poly(I:C) or lipofectamine as a vehicle control (ctrl, n = 18). **e**, Image (top) and quantification (bottom) of Western blot of pSTAT1^Y701^ in ReN VM cell-derived neurons 24h after treatment with deucravacitinib and transfection with poly(I:C) normalized to the housekeeping protein beta actin (*ACTB*). Top: n = 3 per condition. Bottom: n = 9 per condition (all *P* < 0.0001). **f,** Two-way hierarchical clustering of relative protein abundance of IFN-I-related proteins acquired through 10-plex mass spectrometry. Control = lipofectamine (vehicle control). Conditions with drug: n = 2, conditions without drug: n = 3. All replicates in this figure are biological replicates. The error bars represent the standard deviation.

We found interferon receptor alpha and beta subunit 2 (*IFNAR2*; average LFC = 2.23825, *P* = 0.0001), interferon regulatory factor 9 (*IRF9*; average LFC = 2.0625, *P* = 0.0002), and *TYK2* (average LFC = 1.95, *P* = 0.0004) as the best candidates for neural cell rescue upon knockout, whereas knockout of phospholipase C gamma 1 (*PLCG1*; average LFC = -1.9875, *P* = 0.0002), hepatocyte growth factor-regulated tyrosine kinase substrate (*HGS;* average LFC = -1.775, *P* = 0.0004), and *SQSTM1* (p62, average LFC = -1.7475, *P* = 0.0005) markedly increased the toxicity induced by dsRNAmi (Fig. 3b).

We independently validated IFNAR2 and TYK2 using complementary chemical probes. To evaluate IFNAR2, we treated ReN VM cell-derived neural cells with an anti-IFNAR2 antibody before introducing poly(I:C) into the cells (Fig. 3c). The blocking antibody significantly rescued dsRNA-mediated neuronal death, but the response was not dose-dependent and plateaued at about 50% rescue (no antibody *P* < 0.0001; 1:50 *P* = 0.0001; 1:25 *P* = 0.0002; 1:10 *P* = 0.0011). To evaluate our hit TYK2, we used deucravacitinib, a potent small molecule inhibitor ^25^ that is highly specific for TYK2 over other JAK kinase family members (IC_50_ = 0.2 nM compared to IC_50_(JAK1) = 1nM and IC_50_(JAK2) > 200 nM)^43,44^. We observed that exposure to deucravacitinib rescued dsRNA-toxicity up to 100% at all tested concentrations (Fig. 3d) with an EC_50_ < 10 nM. Deucravacitinib was also found to inhibit STAT1 phosphorylation (downstream protein of TYK2) completely at all tested concentrations. (Fig. 3e, all *P* < 0.0001).

To determine whether the activation of IFNAR is sufficient to trigger human neural cell death, we treated cells with interferon-α at a range of doses. After 48 hours, there was no decrease in cell viability at any dose tested (Extended Data Fig. 9e). Taken together, our results indicate that the toxic innate immune reaction to cdsRNA requires the activation of IFNAR, but other factors may play a crucial role in this mechanism leading to neural cell death.

To further evaluate TYK2 as a potential target for inhibiting IFN-I signaling, we performed a deep proteomics analysis from ReN VM cell-derived neural cells that were treated with baricitinib and deucravacitinib at high doses (10 µM) by quantitative 10-plex tandem mass tag (TMT) mass spectrometry (Fig. 3f). Here, we saw that all quantified ISGs were upregulated by poly(I:C), and this ISG induction was reversed by both baricitinib and deucravacitinib.

### Biomarkers for dsRNA pathology

Since no validated biomarkers for TDP-43 inclusions are available, fluid biomarkers associated with dsRNA-induced neuroinflammation in relevant neurodegenerative disease subtypes could help identifying living patients with TDP-43 inclusions. We examined potential biomarkers in cell-based assays by conducting Meso Scale Discovery (MSD) immunoassays on a selected array of ISG products. We quantified CCL2, CXCL10, and IL6 in media taken from ReN VM cells transfected with poly(I:C) and treated with baricitinib, ruxolitinib, deucravacitinib, or vehicle (Fig. 4a-c). The dsRNAmi increased the level of these candidate biomarkers, and the drugs diminished their expression (*P* < 0.0001 in all conditions) to levels comparable to the lipofectamine-only control condition.

**Fig. 4:**
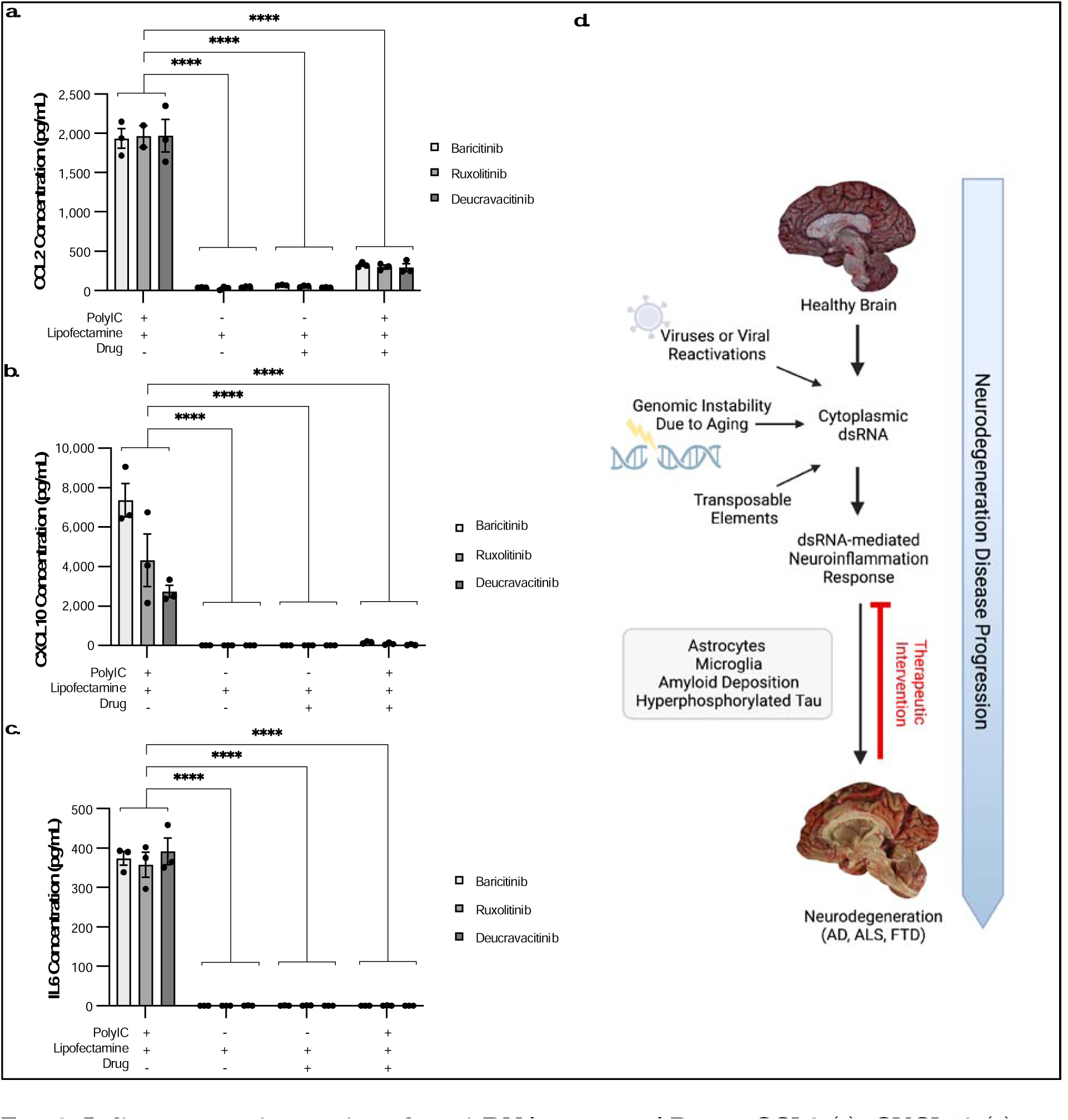
Inflammatory biomarkers for cdsRNA-positive AD. **a-c**, CCL2 (a), CXCL10 (b), and IL6 (c) concentration in pg/mL in the media of ReN VM cell-derived neurons treated with baricitinib, ruxolitinib, or deucravacitinib and transfected with poly(I:C) (n=3 per condition; *P* < 0.0001 in all conditions). **d,** Schematic overview of hypothesized pathomechanisms underlying cdsRNA in neurodegenerative diseases. The error bars represent the standard error of the mean.

## DISCUSSION

Mounting evidence implicates inflammatory signaling as an important driver of AD disease progression, but the root causes of neuroinflammation have been elusive. Here, we demonstrate that cdsRNA, a DAMP, is spatially coincident with cytoplasmic pTDP-43 inclusions in brains of patients with AD and show the upregulation of many ISGs in AD brains which is consistent with the predicted IFN-I response to cdsRNA. We augmented our machine learning pipeline DRIAD-SP^24^ to stratify the cases by expression levels of CEs as a proxy of TDP-43 pathology. Using this new DRIAD-SP pipeline, we identified ruxolitinib and baricitinib as hits in CE-positive subset of AD, further refining the hypothesis of JAK inhibitors as drug candidates to a subset of AD patients with TDP-43 dysfunction. Both our CRISPR screen and systems pharmacology analyses implicated the selective inhibition of TYK2 as a target of rescuing cdsRNA-induced toxicity. Our findings are consistent with previous studies (including our own^24^) linking TYK2 to AD^45–47^. Finally, we propose CCL2, CXCL10 and IL6 as candidate inflammatory biomarkers in neurodegenerative diseases with cdsRNA-pathology.

The accumulation of cdsRNA in neurons with pTDP43 dysfunction^48^ may be an unappreciated pathogenic mechanism in AD as a root cause of neuroinflammatory cascades that are triggered within neurons. Additional evidence supporting this concept of “neuroninflammation” includes expression of cdsRNA exclusively in neurons. Introducing dsRNA is sufficient to trigger neuroinflammation and neuronal death in the Nd1 and Nd2 mouse models^4^ and cdsRNA also induces an innate immune response and death in a differentiated human neural cell-based model that lacks microglia. However, growing evidence supports microglia activation playing an important role in AD progression, and cdsRNA-mediated neuroinflammation and neural cell toxicity can trigger microglial activation^4,48,49^ and consequent propagation of AD pathology^1,2^.

In a tau-based mouse model, mitochondrial DNA leakage leading to cytoplasmic DNA and STING activation in microglia could contribute to an IFN-I pathogenic mechanism^50^. Additionally, cytoplasmic DNA within neurons could activate this cGAS-STING pathway ^51^ which activates the kinase TBK1, a common node of dsRNA-and dsDNA-mediated innate immune signaling networks.

Drug repurposing accelerates testing therapeutic hypotheses expeditiously in clinical trials, and FDA approved drugs can serve as chemical biology probes to elucidate novel targets. By elucidating the mechanism of action of baricitinib and ruxolitinib in human neural cells, we identified TYK2 as a target to rescue cdsRNA-induced toxicity. Deucravacitinib reverses dsRNA-induced toxicity at lower doses than either ruxolitinib or baricitinib. Its high selectivity for TYK2 could achieve improved safety profiles relative to baricitinib and ruxolitinib. The inhibition of JAK1-3 is associated with adverse infectious, embolic, and thrombotic, neoplastic, and gastrointestinal perforation events^52^, which have not been reported in people with TYK2 polymorphisms^53,54^ or those treated with a potent TYK2 inhibitor^55^. However, deucravacitinib is unlikely to cross the blood brain barrier based on its polar molecular structure^56^, making it imperative to develop novel brain-penetrant TYK2-selective inhibitors.

Translation of these results will require identification of validated predictive biomarkers of drug action to the inflammatory response. We have identified candidate biomarkers to detect the activation of cdsRNA-mediated neurodegeneration in the CSF of patients which affords the possibility of precision medicine in clinical trials for AD and other neurodegenerative diseases. In addition to our proposed biomarkers, recent studies show that peptides expressed from RNA containing CEs as a result of nuclear TDP-43-depletion can be detected in the CSF of ALS patients^57,58^ and could thus also be used as a biomarker in AD and other cdsRNA/TDP-43-positive neurodegenerative diseases, such as chronic traumatic encephalopathy^59^. We are conducting a clinical trial in AD patients (NCT05040659) to quantify the blood brain barrier penetrance of baricitinib and to evaluate the response of inflammatory and neuronal death biomarkers.

Limitations to our study include that we do not observe spatially coincident accumulation of cdsRNA foci and NFTs, which might be expected based on recent work illustrating the connection between tau pathology and transposon derepression^60^. One reason for this could be that the toxicity of spatially coincident cdsRNA and NFTs in neurons in the examined brain regions is too high to be able to observe cells expressing both pathologic lesions in autopsy samples. Our analysis included a limited number of brain regions both in our histologic and our computational analyses. Future studies will analyze more brain regions in cohorts of more diverse patients. Also remaining to be determined is the source of cdsRNA in AD. We hypothesize several potential sources of cdsRNA (Fig. 4d), including derepression of transposable elements^61^ such as inverted Alu elements^21,62^, genomic instability that occurs naturally in an aging brain, and the reactivation of viruses^63^ such as HSV^64^. Future studies will deploy computational methods (quantitative trait locus analysis) and the isolation and sequencing of cdsRNA from autopsied brains from sporadic AD cases.

In summary, we hypothesize that cdsRNA-induced innate immune responses contribute to disease progression in the subset of AD patients with TDP-43 pathology and also in other neurodegenerative diseases with TDP-43 inclusions, including ALS and FTD (Fig. 4d). This cdsRNA may accumulate over the years until it reaches a triggering point as a DAMP where it activates PRRs that activate IFN-I signaling and neuroinflammation. This inflammatory pathology is intertwined with amyloid deposition and the hyperphosphorylation of tau, either as a parallel pathomechanism or placed upstream or downstream of known proteinopathies. The involvement of astrocytes and microglia in dsRNA-induced toxicity has been seen in our mouse models^4^ and others^48^, but the data from our human cell-based models suggest that microglia are not necessary for neural cell death. Therapeutic interventions that target TYK2 to inhibit cdsRNA-induced neuroinflammation may have potential not only in AD but also in other neurodegenerative diseases associated with TDP-43 pathology.

## Supporting information

SI Guide

## METHODS

### Histological Staining and Imaging

All human tissues were acquired from the MGH MADRC brain bank in accordance with protocols approved by the MGB Institutional Review Board. Similar to what was previously reported^4^, ∼5 µm thick formalin-fixed paraffin-embedded (FFPE) sections of the amygdala were dewaxed followed by antigen retrieval in a Leica Bond Fully Automated Slide Stainer. After deparaffination and rehydration by sequential incubation in Leica dewing solution (3 × 5 min), 100% ethanol (2 × 5 min), 90% ethanol (2 × 5 min), 70% ethanol (2 × 5 min), and distilled water (3 × 5 min), antigen retrieval was achieved with citrate-based antigen retrieval solution (ER2) and steam cooking for 1 hour. Afterwards, sections were cooled to room temperature and rinsed in water and 1× PBS. For permeabilization, 0.03% hydrogen peroxide solution was applied for 8 min. After blocking in tris-buffered saline (TBS)–based Odyssey blocking buffer (cat. # 927-60001, LI-COR, NE, USA) for 1 hour, for immunohistochemistry, primary antibodies (J2 for dsRNA (cat. # 76651, Cell Signaling Technology, MA, USA); pTDP-43 S409/410 (cat. # CAC-TIP-PTD-P02, Cosmo Bio, CA, USA); tau (cat. # NBP2-25162, Novus Biologicals, CO, USA); Aß (cat. # CST-D54D2, Cell Signaling Technology)) diluted in blocking buffer were applied overnight at 4°C in a humidified chamber. Following a wash with PBS (3 × 5 min), secondary antibodies were applied using the VECTASTAIN ABC Staining Kit (cat. # PK-6200, Vector Laboratories, CA, USA) and DAB Staining Kit (cat. # SK-4100, Vector Laboratories). For luxol fast blue-hematoxylin and eosin (LHE) staining, the protocol of the Luxol Fast Blue Stain Kit (cat. # ab150675, abcam, MA, USA) and the Sakura Prisma H&E Stain Kit (cat. # 76318-76, VWR, PA, USA) were followed. For tissue-based cyclic immunofluorescence (CyCIF)^27^, unspecific binding sites were blocked, slides were incubated in 3% H_2_O_2_ and 20 mM HCl in PBS at room temperature with light-emitting diode illumination for 1 hour to reduce autofluorescence, and the first primary antibody set (J2 and pTDP-43 S409/410) was applied. After washing, the secondary antibodies (anti-rabbit Alexa Flour 555 (cat. # A-31572, Thermo Fisher, MA, USA); anti-mouse Alexa Flour 647 (cat. # A-21237, Thermo Fisher)) 1:1000 diluted in blocking buffer were applied for 1 hour in a humidified chamber at room temperature. After a wash, Sytox Blue Nucleic Acid Stain (cat. # S11348, Thermo Fisher) was applied 1:15000 in blocking buffer for 15 min at room temperature. Following a final washing step, slides were imaged. After this first cycle, the fluorophores were inactivated by 4.5% H_2_O_2_ and 24 mM NaOH in PBS at room temperature under light exposure for 1 hour. After rinsing, the second primary antibody set (tau and Aß) was applied and staining was continued in the same manner as described above.

### Differential Gene Expression Analysis of Different Brain Regions

Differential Gene Expression was conducted based on a previously established protocol^65^. In brief, clinical diagnosis was used to define AD versus control conditions within the bulk RNA-sequencing data taken from the ROSMAP^29^ and MSBB^30^ databases (data obtained from the AMP-AD Knowledge Portal (doi:10.7303/ syn2580853)). The analysis focused on a list of 354 pre-selected interferon-related genes. Of these genes, 309 could be found in the analyzed brain regions (from ROSMAP: dorsolateral prefrontal cortex (DFPC), from MSBB: frontal pole (FP), inferior frontal gyrus (IFG), parahippocampal gyrus (PHG), superior temporal gyrus (STG)). 139 genes were significantly upregulated (adjusted *P* value < 0.05) while 51 were downregulated. The full results with logFC, FC, CI, *P* values and adjusted *P* values can be found in Supplementary Table 2.

### ReNcell VM Culturing and Testing

ReN VM cells (cat. # SCC008, Millipore, MA, USA) were cultured in ReNcell Maintenance media (cat. # SCM005, Millipore) containing 20 ng/ml of epidermal growth factor (cat. # GF001, Millipore) and 20 ng/ml of basic fibroblast growth factor (cat. # 03-002, Stemgent, MD, USA) and differentiated into neural cells for one week by removing the growth factors as previously described^4,66^, with the difference that the differentiation took place in a separate dish (Extended Figure 8) to minimize variability downstream. Differentiated cells were seeded into a Matrigel-coated (cat. # 354320, Corning, NY, USA) 96-well plates (20,000 cells per well) and where applicable, respective drugs dissolved in dimethyl sulfoxide (DMSO) (baricitinib: HMS LINCS ID # 10354, ruxolitinib: HMS LINCS ID # 10138, deucravacitinib: cat. # HY-117287, MedChem Express, NJ, USA) were added using a D300e Digital Dispenser (Hewlett-Packard, CA, USA). After 1 hour of incubation at 37 °C, cells were transfected with polyIC HMW (cat. # tlrl-pic, Invitrogen, CA, USA) by lipofection with lipofectamine 2000 (cat. # 11668027, Thermo Fisher) (final dsRNA concentration: 8 µg/ml). To test the relevance of IFNAR2, a neutralizing anti-IFNAR2 antibody (cat. # 21385-1, PBL Assay Science, NJ, USA) was added in the respective amount to each well after lipofection and without prior drug treatment. To test whether interferon α and polyIC have comparable effects on neurons, interferon α (cat. # 407294-5MU, Millipore) was diluted in media and added to the cells without prior drug treatment. Western blot samples and cell media for biomarker testing were collected 24h post-treatment. Cell viability was assessed 72 hours post-treatment using the CellTiter-Glo Assay (cat. # G7572, Promega, WI, USA) following the manufacturer’s instructions. All conditions were normalized to 0.1% DMSO content. Where applicable, lipofectamine 2000 was used as control. For mass spectrometry, two million ReN VM cells were expended in a 15-cm-dish (2 dishes per replicate) for 4 days and subsequently differentiated for 1 week, then treated with 10µM of drug, transfected with polyIC after 1 hour of incubation at 37 °C, and 24 hours later, washed with ice-cold 1x PBS and collected by scraping followed by centrifugation at 500 g for 5 min.

### RNA-sequencing data processing and prediction of TDP-43 status

RNA-sequencing reads of brain specimens from Alzheimer’s disease patients in the ROSMAP study, along with their corresponding clinical annotations, were downloaded from the AMP-AD Synapse portal at https://adknowledgeportal.synapse.org. RNA-sequencing reads from ALS patient neurons that were sorted for presence or absence of TDP-43 were downloaded from Sequence Read Archive (SRA) accession GSE126542^36^. We quantified transcripts using Salmon v1.9.0^67^ against release 107 of the human transcriptome from Ensembl. We amended the transcriptome with three transcripts that have not yet been annotated in Ensembl, but that are known to be associated with loss of nuclear TDP-43^17^. Specifically, we included two splice variants of UNC13A, each including an additional CE between the canonical exons 20 and 21 (hg38; chr19: 17,642,414–17,642,541 (CE1); chr19: 17,642,414–17,642,591 (CE2)), and one splice variant of STMN2, which includes an alternative exon 2 termed exon 2a (hg38; chr8:79,616,822–79,617,048).

Given our quantification of TDP-43 pathology associated transcripts we classified patient samples as TDP-43-positive or -negative based on the number of expressed CEs. We dichotomized transcript abundance using thresholds set above the lowest non-zero peak in the abundance histograms (TPM>1.3 STMN2short; TPM>0.09 for UNC13A-CE1; TPM>0.08 UNC13-CE2). This threshold determines whether each individual CE is present or absent. Next, patient samples expressing one or none of the three CEs (STMN short, UNC13A-CE1 or UNC13A-CE2) were considered TDP-43-negative and patient samples expressing two or all three of the CEs were considered TDP-43-positive. Using this classification, the proportion of TDP-43 pathology positive samples is approximately 37%. Our classification is a conservative prediction, given that the clinical incidence of TDP-43 inclusions in AD patients is up to 57%^13^.

### Prediction of disease stage and assessing gene set significance

For assessing how well any given gene set predicts Alzheimer’s disease progression we used an updated variant of the published DRIAD-SP methodology^24^. For predicting disease stage, we collapsed transcript abundances to gene-level abundances using tximport^68^. The entire transcriptional feature space was filtered down to the about ∼20k protein-coding genes in the human genome to ensure that only genes with gene products that can conceivably be targeted using conventional small molecule inhibitors are included. Every AMP-AD specimen was assigned a label of disease severity based on the following mapping to the Braak annotations: A—early (Braak 1–2), B—intermediate (Braak 3–4), and C—late (Braak 5–6). We set up an ordinal regression task classifying patients according to their disease stage using the ordinalRidge R package (https://github.com/labsyspharm/ordinalRidge). As predictors we used the gene expression matrix from AMP-AD, filtered to only contain genes in the current gene set under investigation. This way, the performance (AUC) of a model utilizing a given gene set can be viewed as a measure of how strongly the given genes are predictive of AD progression. To address overfitting, ordinalRidge utilizes a ridge regularization term that penalizes the L2-norm of feature weights. No LASSO regularization was used, as it induces sparsity and excludes features that were specifically preselected to be included in the model.

Model performance was evaluated through leave-pair-out cross-validation. For a given regression task, each example in the dataset was associated with an example from the other classes that was the closest match in age. If there were multiple candidates for the age match, the pairing was selected uniformly at random. The resulting set of age-matched pairs was evaluated in a standard cross validation setting, by asking whether the later-stage example in each withheld pair was correctly assigned a higher score by the corresponding predictor. The fraction of correctly ranked pairs constitutes an estimate of the area under the ROC curve^69^.

Comparing model performances across gene sets directly is not possible, as their set sizes can differ. Therefore, we normalized AUC values by relating them to performances observed on size-matched random gene sets. For each gene set of interest, 1000 random gene sets of matching sizes were sampled from a uniform distribution over the protein-coding space. After evaluating all lists through cross-validation, an empirical *P* value was computed as the fraction of background sets that yield higher predictor performance than the gene list of interest.

### CRISPR screen

Low passage (P2) ReNcell VM cells stably expressing Cas9 were expanded and transduced with lentiviral particles containing the Brunello library^42^. Lentiviral transduction efficiency was tested by plating cells into two 6-well plates and varying volumes of stock preparation of Brunello library lentiviral vectors with 8 µg/ml of polybrene. One plate was treated with puromycin, the other plate remained untreated, and the proportions of cells measured for each condition using CellTiter-Glo were used to calculate the efficiency which was found to be at 84% averaged from triplicates. Transduced cells were expanded in the presence of puromycin until the required number of cells was achieved to obtain triplicates of 1000 cells per sgRNA. Then, cells were differentiated as described above. After seven days, 3 µg/ml dsRNA in complex with lipofectamine (concentration according to manufacturer’s protocol) or lipofectamine alone as a control were added to each flask. After 3 days of treatment, genomic DNA was isolated from each group of cells. PCR and sequencing were performed as previously described^70,71^. For analysis, the read counts were normalized to reads per million and then log2 transformed and the LFC of treatment and control were compared to the lipofectamine controls.

### Mass spectrometry

First, cells were lysed in lysis buffer (2% SDS, 150 mM NaCl, 50 mM Tris pH 7.4) supplemented with protease and phosphatase inhibitor (cat. # 78445, Thermo Fisher) and homogenized with a QIAshredder column (cat. # 79656, Thermo Fisher). Lysates were reduced with freshly prepared dithiothreitol (DTT) (final concentration: 5 mM) and heated at 37 °C for 1 hour. Samples were then alkylated with iodoacetamide (final concentration:15 mM) for 30 min in the dark whereupon the reaction was stopped by adding DTT (final concentration: 50 mM).

After methanol/chloroform precipitation, the purified protein was solubilized in freshly prepared 8 M urea in 20 mM EPPS, at pH 8.5 for 30 mins at 37 °C. For digestion, the urea concentration was diluted with 20 mM EPPS to 4 M final concentration and 2% acetonitrile (v/v) was added. Digestion was performed using lysyl endopeptidase® Lys-C (2 mg/ml; cat. # 121-05063; Wako, Osaka, Japan), (enzyme-to-substrate ratio: 1:50) for 3 hours at 37 °C. Following dilution of the sample with 20 mM EPPS to a final urea concentration of 1.6 M, further digestion was performed by the addition of trypsin (cat. # V5111, Promega), (enzyme-to-substrate ratio: 1:100) for 8 hours at 37 °C. A small aliquot was analyzed for its missed cleavage rate by mass spectrometry. Equal amounts of digested protein (60 µg) were removed from each sample and labeled using TMTpro 16plex Mass Tag Labelling Kit (TMT; cat. # A44520, Thermo Fisher) following the manufacturer’s instructions. Labeling efficiency of > 95% was determined along with ratio checks by mass spectrometry MS^3^, while the labeling reactions were stored at -80 °C. To quench the reaction, hydroxylamine was added to a final concentration of 0.5% (v/v) for 10 mins. After acidification using formic acid, equal amounts of labelled peptide were pooled from each sample (as judged from ratio check data). Then, solvent was evaporated. The labeled peptide multiplex was desalted by solid-phase extraction (SPE; SepPak tC18 Vac RC Cartridge). The peptides were fractionated by HPLC alkaline reverse phase chromatography (Agilent 1200 Series) into 96 fractions and combined into 24 samples. Lastly, peptides were desalted over Stage Tips^72^ prior to MS analysis.

A MultiNotch SPS-MS^3^ TMT method^73^ was used on an Orbitrap Lumos mass spectrometer (Thermo Fisher) coupled to a Proxeon EASY-nLC 1200 liquid chromatography (LC) system (Thermo Fisher). Samples were injected onto a 40 cm, 100 µm (internal diameter) column packed with 2.6 µm Accucore C^18^ resin (flow rate of 450 nl/min). Over the course of 4 hours, the peptide fractions were separated through acidic acetonitrile gradients by the LC before being injected into the mass spectrometer. First, an MS^1^ spectrum (Orbitrap analysis; resolution 120,00; mass range 400-1400 Th) was taken. Then, an MS^2^ spectrum was collected after collision-induced dissociation (CID, CE = 35) with a maximum ion injection time of 150 ms and an isolation window of 0.7 Da. For TMT quantification of peptides, MS^3^ precursors were fragmented by high-energy collision-induced dissociation (HCD, CE = 40%) and analyzed in the Orbitrap at a resolution of 50,000 at 200 Th. Further details can be found in a previously published article^74^.

A Sequest-based in-house software was used to search peptides against a human database with a target decoy database strategy and a false discovery rate of 2%. Oxidized methionine residues (+15.9949 Da) were dynamically searched, along with static modifications for alkylated cysteines (+57.0215 Da) and the TMTpro reagents (+229.1629 Da) on lysines and the N-termini of peptides. Relative protein quantification required a summed MS^3^ TMT signal/noise > 200 over all TMT channels per peptide and an isolation specificity > 70% for any given peptide. Quant tables were generated and exported to Excel for further procession. More details on the TMT intensity quantification and certain parameters can be found in another recent publication^75^.

### Western blotting

First, 15 µl of x Laemmli Sample Buffer (cat. # 1610737; Bio-Rad Laboratories, CA, USA) with 1:20 2-mercaptoethanol was added to each well after aspiration of the media. Subsequently, the lysates were boiled at 100 °C for 5 min. Then, the samples were loaded onto a precast gel (cat. # 4569036; Bio-Rad Laboratories, CA, USA) which was run in 1x running buffer diluted from a 10x Tris/Glycine/SDS stock (cat. # 1610772; Bio-Rad Laboratories) at 160 V for 40 min. The transfer was performed using a PVDF membrane (cat. # IPFL00010; Millipore) and a 1x transfer buffer diluted from a 10x stock (cat. # PI35040; Thermo Fisher, MA, USA) mixed with 20% methanol. For 90 min, the gel was blotted at 4 °C at 90 V. Finally, unspecific binding sites were blocked by phosphate-buffered saline (PBS)-based Odyssey® blocking buffer (cat. # 927-40150; LI-COR) for 1 hour at room temperature. Then, primary antibody (pSTAT1 Y701 (cat. # 9167S; Cell Signaling Technology); beta actin (ACTB, cat. # 3700; Cell Signaling Technology) diluted 1:1000 in blocking buffer was applied overnight at 4 °C. Then, the blot was washed with tris-buffered saline (cat. # sc-262305; Santa Cruz, TX, USA) containing 0.05% Tween 20 (cat. # BP337-500; Thermo Fisher) (3 x 5min). Afterwards, secondary antibody (IRDye 800CW (cat. # 926-32211; LI-COR); IRDye 680RD (cat. # 926-68070; LI-COR)) diluted 1:2500 in blocking buffer was applied for 1 hour at room temperature. After three final washes, the blot was imaged with the Odyssey® DLx Imager (LI-COR). Images were analyzed using ImageStudioLite (LI-COR).

### Meso Scale Diagnostics (MSD) biomarker assay

To measure CCL2 (MCP-1), CXCL10 (IP-10), and IL-6 biomarker levels in collected cell media, we used commercially available V-PLEX assays from MSD, LLC (Cambridge, MA, USA). For all assays, we used the proposed procedures with the exception that different dilutions were required for the sample values to fall within the dynamic range of the assay. For V-PLEX MCP-1 Human Kit (cat. # K151NND), samples were diluted 1:1000 to fit within the 0.09-375 pg/mL range. For V-PLEX IP-10 Human Kit (cat. # K151NVD) and V-PLEX IL-6 Human Kit (cat. # K151QXD), samples were diluted 1:100 to fit within the 0.37-500 pg/mL and 0.06-488 pg/mL ranges, respectively. All plates were analyzed under the MESO QuickPlex SQ 120 (Model # 1300) plate reader and concentrations were quantified via the Meso Scale Discovery WorkBench. Samples with concentrations reported as “NaN” were quantified as zero, as these concentrations fell below the dynamic ranges reported above.

### Statistical Analysis

Statistical analyses were conducted on GraphPad Prism for unpaired *t* tests with two tails and ordinary one-way analysis of variance (ANOVA) with post hoc tests. Correction for multiple comparison was done using Dunnett’s statistical hypothesis testing. Differences were considered significant if *P* < 0.05. P value style: *P* > 0.05 and *P* = 0.05 (no asterisks), *P* < 0.05 (*), *P* < 0.01 (**), *P* < 0.001 (***), *P* < 0.0001 (****). No outliers were eliminated from the dataset.

## DATA AVAILABILITY

For all compounds with an HMS LINCS ID, the compound information, including the vendors they were purchased from, can be found on the HMS LINCS website (https://lincs.hms.harvard.edu/). The proteomics raw data and search results have been deposited to the ProteomeXchange Consortium via the PRIDE^76^ partner repository with the dataset identifier PXD043641 (note to reviewers: datasets will be publicly available when paper is published, per PRIDE policy; data are available during the review phases at https://www.ebi.ac.uk/pride/login with username, password: AYavnzhM).

## CODE AVAILABILITY

Scripts to fully reproduce the tables and figures represented in this manuscript are provided on GitHub at https://github.com/labsyspharm/ad-personalized.

## ACKNOWLEDGEMENTS

We thank Kyle Evans for technical support; and Brad Hyman, and Steven Arnold for sharing equipment and insightful discussions. We thank R01 AG058063 (awarded to M.W.A. and P.K.S.); RF1 AG078297) (awarded to M.W.A.), U54CA225088 (subaward to M.W.A. of overall award to P.K.S.), QFASTR award from MGH (awarded to M.W.A.), the CART fund (awarded to M.W.A.), the MassCATS program (awarded to M.W.A.) for support.

The results published here are in part based on data obtained from the AMP-AD Knowledge Portal (doi:10.7303/ syn2580853). These data were generated from postmortem brain tissue collected through the Mount Sinai VA Medical Center Brain Bank, led by Dr. Eric Schadt from Mount Sinai School of Medicine, and through the Rush Alzheimer’s Disease Center, led by Dr. David Bennett from the Rush University Medical Center, Chicago.

## AUTHOR CONTRIBUTIONS

L.E.K., S.R. and M.W.A. designed the experiments. L.E.K., S.R., Sh.D., A.C., A.S. and G.Z. contributed to the investigative work, data curation, and the validation of the presented findings.

C.H. and A.S. implemented the machine learning framework and conducted the drug prediction analysis. Su.D. contributed to the computational analysis of RNA-sequencing data regarding interferon signaling. G.B., R.E. and M.K. collected, analyzed, and visualized the proteomics data. F.P. and D.R. contributed to the CRISPR screen. L.E.K. and M.W.A. wrote the original draft of the manuscript. L.E.K. and Sh.D. contributed to the visualization of the results. All authors contributed to figures and manuscript reviewing and editing. P.S. contributed to the conceptualization and supervision of this project. M.W.A. was responsible for supervision, project administration, and funding acquisition.

## COMPETING INTERESTS

M.W.A. is a consultant for TLL, LLC, Transposon Therapeutics, and has received in kind support from Eli Lilly that is not related to this work. A.S. is an employee at Flagship Labs 84, Inc., a subsidiary of Flagship Pioneering. F.P. is an employee of Merck Research Laboratories.

## ADDITIONAL INFORMATION

Supplementary Information is available for this paper.

Correspondence and requests for materials should be addressed to Mark W. Albers.

Reprints and permissions information is available at www.nature.com/reprints.

## EXTENDED DATA

**Extended Data Table 1:** Patient data of human postmortem brains sections.

| Patient information |  |  |  |  |  |  |  |  |
| --- | --- | --- | --- | --- | --- | --- | --- | --- |
| MADRC Number | Age at death | Sex | Race | PMI [hours] | NPDX | CERAD | Braak staging | Figure reference |
| 1641 | 63 | male | white | 16 | healthy control | NA | NA | Figure 1a (Healthy Control Case I) |
| 1747 | 65 | male | white | NA | healthy control | NA | NA | Figure 1a (Healthy Control Case II), Figure 1b (IHC of Healthy Control), Extended Data Figure 1b (Normal) |
| 2009 | 94 | male | NA | 23 | healthy control | A possible | II | Figure 1a (Healthy Control Case III), Figure 1b (IF of Healthy Control), Extended Data Figure 1c (Control), Extended Data Figure 2 (Control) |
| 1828 | 91 | female | white | 16 | Alzheimer's Disease | B probable | VI | Figure 1a (Alzheimer's Disease Case I), Figure 1b (IHC of Alzheimer's Disease), Extended Data Figure 1b (Severe) |
| 1896 | 82 | male | white | 16 | Alzheimer's Disease | B probable | VI | Figure 1a (Alzheimer's Disease Case II), Figure 1b (IF of Alzheimer's Disease), Extended Data Figure 1b (Mild) and 1c (Alzheimer's Disease), Extended Data Figure 2 (Alzheimer's Disease - left) |
| 1966 | 84 | female | white | NA | Alzheimer's Disease | C definite | VI | Figure 1a (Alzheimer's Disease Case III) |
| 1967 | 70 | male | NA | 6 | Alzheimer's Disease | C definite | V | Extended Data Figure 2 (Alzheimer's Disease - middle) |
| MADRC = Massachusetts Alzheimer's Disease Research Center<br>NPDX = Neuropathology diagnosis<br>PMI = Postmortem Interval<br>CERAD = The Consortium to Establish a Registry for Alzheimer's Disease |  |  |  |  |  |  |  |  |

**Extended Data Fig. 1:**
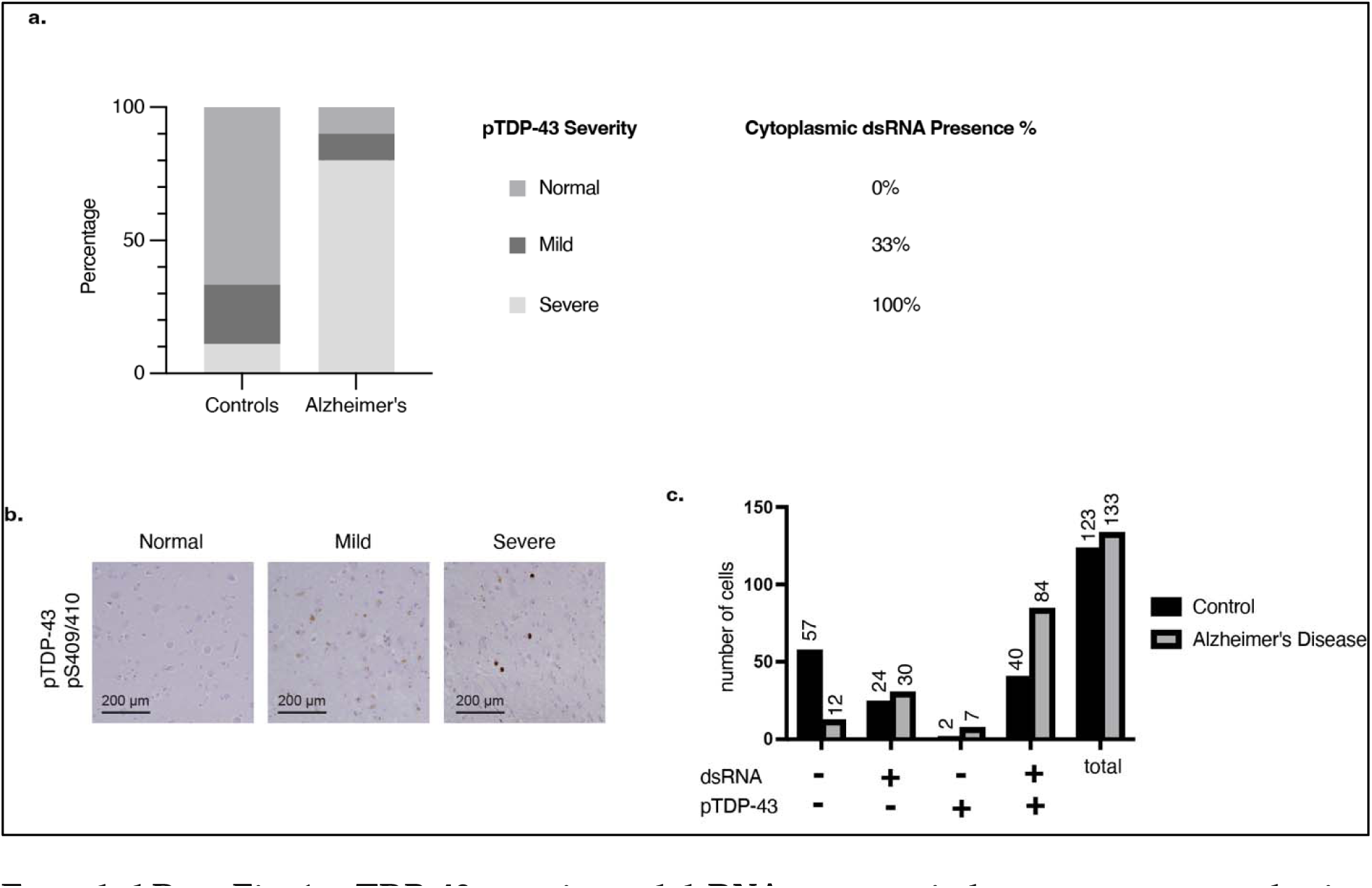
pTDP-43-severity and dsRNA presence in human postmortem brain sections. **a**, Percentage of assessed FFPE sections (n = 10 with AD, n = 9 healthy controls) that show normal, mild, and severe pTDP-43 severity compared to healthy controls as well as the coincidence with dsRNA in percent of cases. **b,** Histological examples for the classification of pTDP-43 severity into normal, mild, and severe. **c,** Proportion of cells that stained for cdsRNA, pTDP-43, or both.

**Extended Data Fig. 2:**
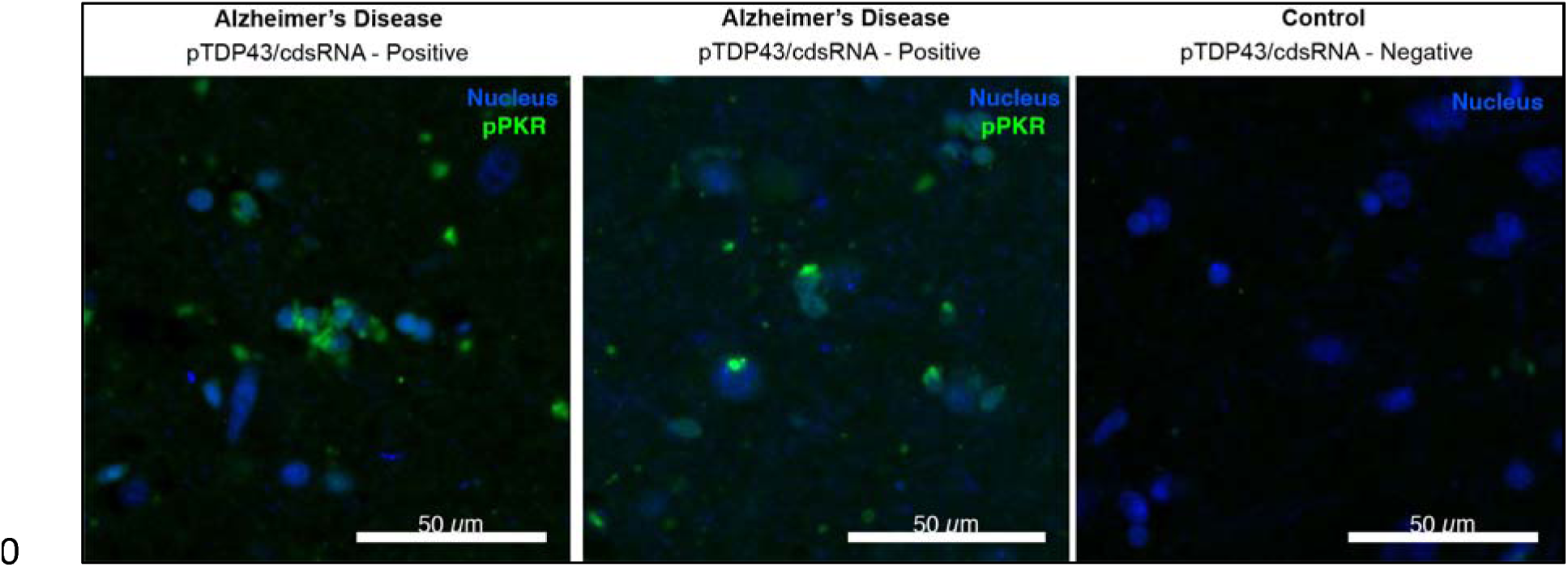
Phosphorylation of PKR as proof of the immunogenicity of cdsRNA. Immunofluorescence staining of human postmortem brain sections of the amygdala comparing the phosphorylation of PKR (green) in pTDP-43/cdsRNA-positive AD (left and middle image) to a pTDP-43/cdsRNA-negative control (right image). Sytox Blue used to stain nuclei.

**Extended Data Fig. 3:**
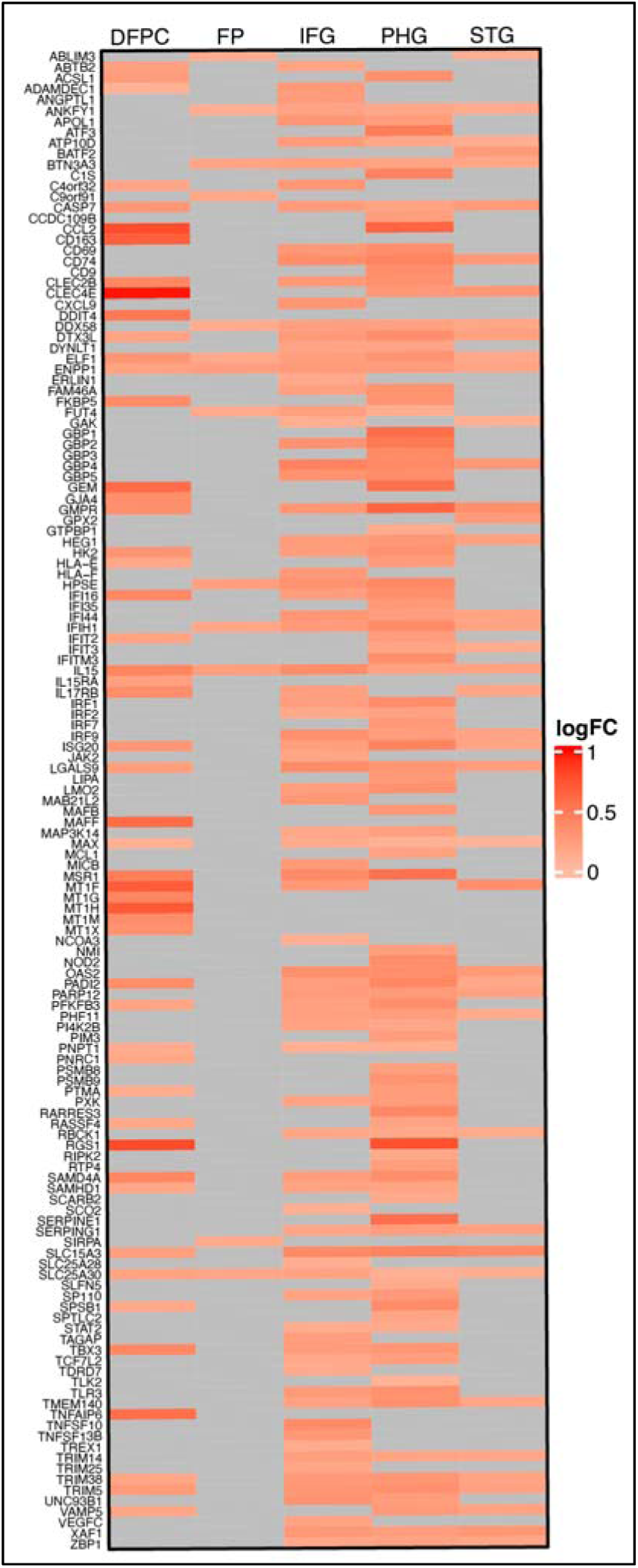
Upregulated ISGs within brain regions relevant in AD. Heat map of RNA-sequencing data derived from the ROSMAP^29^ and MSBB^30^ databases comparing AD patients and healthy controls. Differential expression shown in log2 fold change (logFC) for pre-selected ISGs. DFPC: dorsolateral prefrontal cortex; FP: frontal pole; IFG: inferior frontal gyrus; PHG: parahippocampal gyrus; STG: superior temporal gyrus.

**Extended Data Fig. 4:**
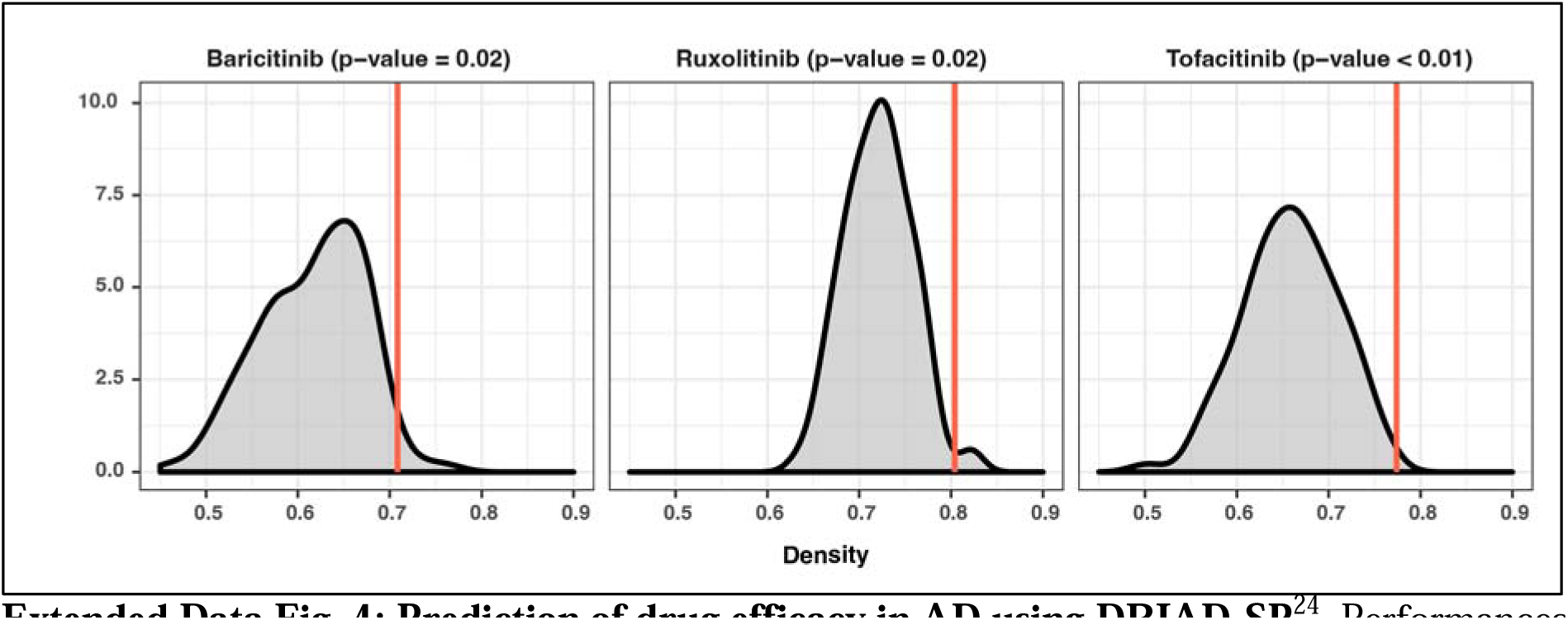
Prediction of drug efficacy in AD using DRIAD-SP^24^. Performances of three selected drugs in the DRIAD-SP prediction task. Drug efficacy was assessed according to the previously published protocol^24^. Red lines indicate the DRIAD-SP model performance of the drug gene sets, whereas the gray shaded regions correspond to the distribution of model performances based on size-matched random gene sets.

**Extended Data Fig. 5:**
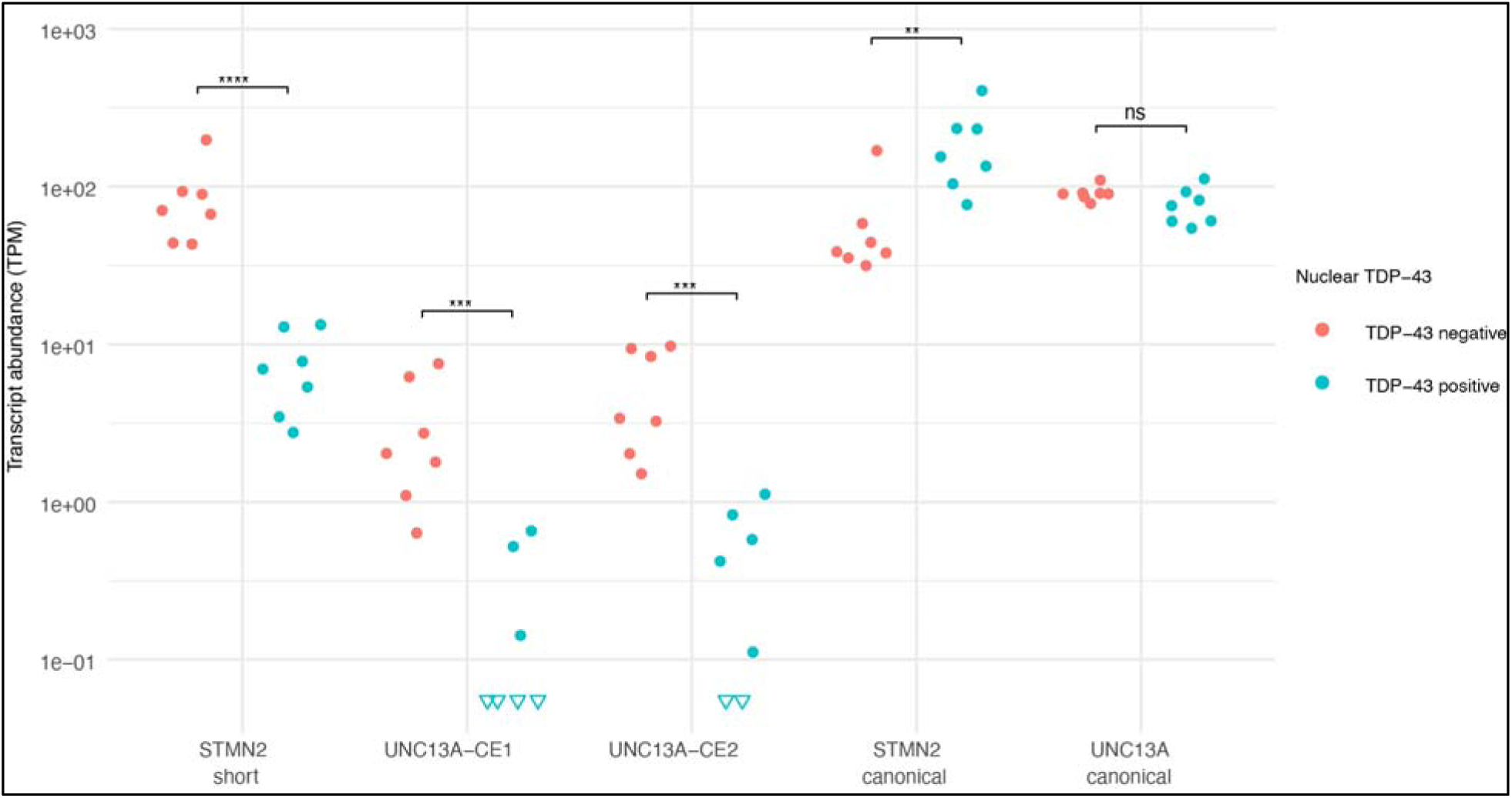
Expression of TDP-43-associated transcripts in single neuronal nuclei. Neuronal nuclei from ALS patients’ neocortical tissues were FACS-sorted by the presence of nuclear TDP-43 and NeuN^36^. Here, we sorted the RNA-sequencing data obtained from the sorted nuclei for the presence of CEs as a proxy of TDP-43 pathology and thus absence of nuclear TDP-43. Each point corresponds to the abundance (transcripts per million; TPM) of TDP-43-associated CE transcripts or their canonical counterparts in one of the samples. (STMN2 short *P* < 0.0001; UNC13A-CE1 *P* = 0.0002; UNC13A-CE2 *P* = 0.0006; STMN2 *P* = 0.0017; UNC13A *P* = 0.11).

**Extended Data Fig. 6:**
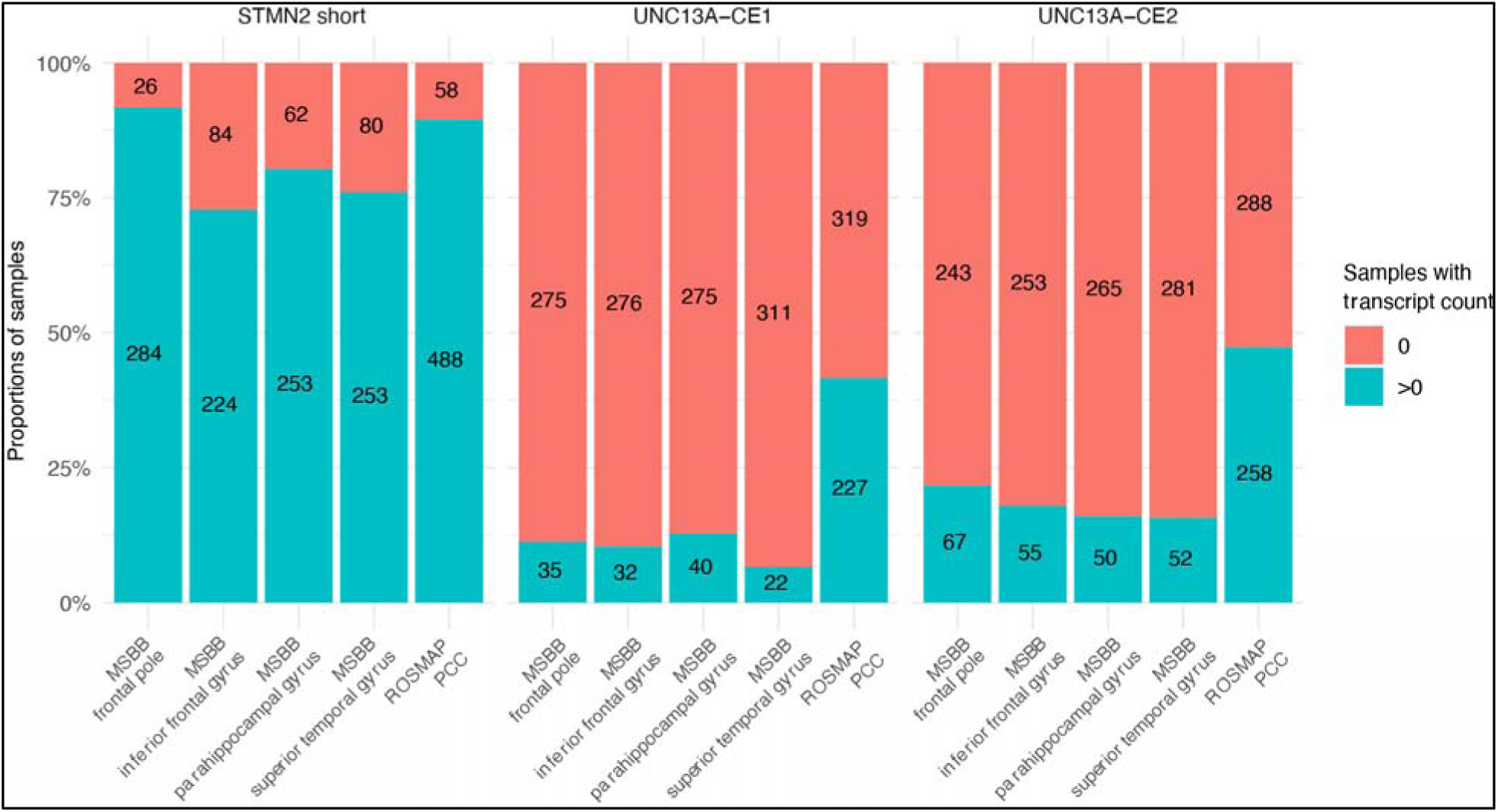
Comparison of CE transcript abundance between ROSMAP^29^ and MSBB^30^ datasets. The proportion of samples with zero (red) and above zero (blue) abundance (transcripts per million; TPM) of the given transcripts is shown. MSBB samples had an appreciable lower abundance of UNC13A CE transcripts compared to ROSMAP samples. This difference can most likely be attributed to MSBB using single-end sequencing compared to the paired-end sequencing employed by ROSMAP, making determination of TDP-43 pathology in MSBB difficult.

**Extended Data Fig. 7:**
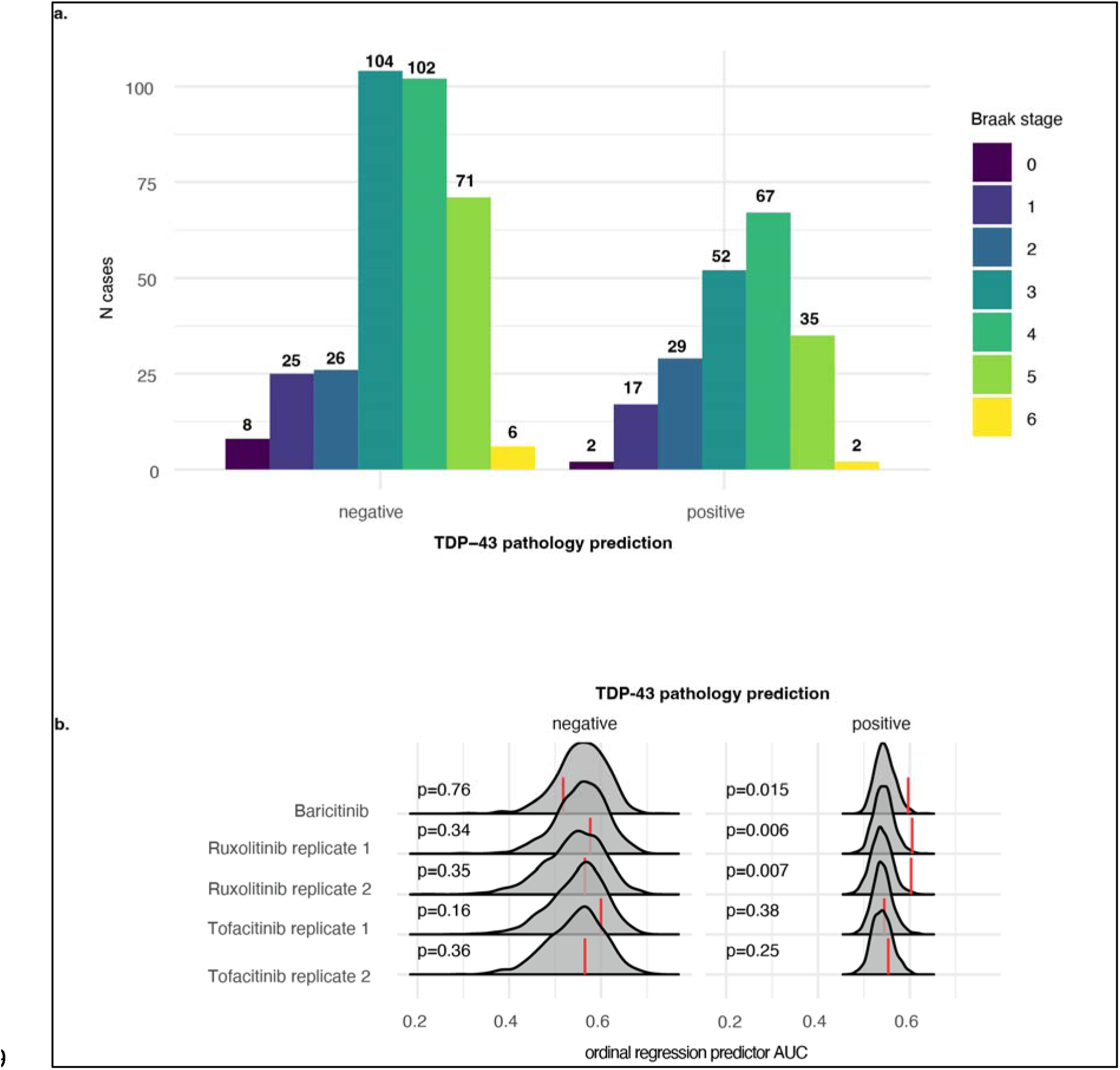
Braak staging of TDP-43 cases and additional replicates for the prediction of drug efficacy through DRIAD-SP. **a,** Comparison of Braak stage distribution of ROSMAP patient data (posterior cingulate cortex) relative to their predicted TDP-43 pathology. **b,** Performance of baricitinib, ruxolitinib and tofacitinib in DRIAD-SP as shown in Figure 2e, here including additional replicates of ruxolitinib and tofacitinib.

**Extended Data Fig. 8:**
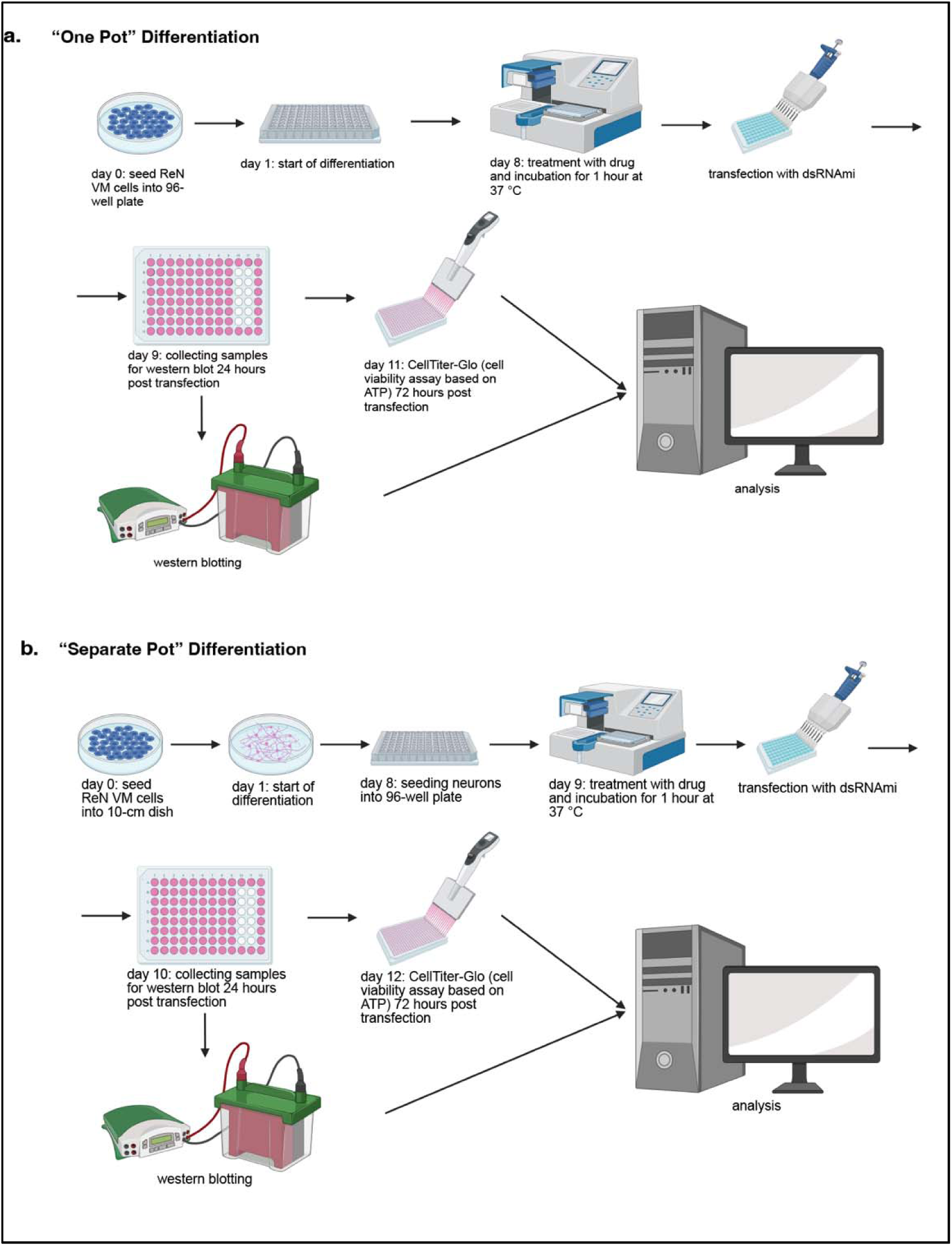
Schematic overview comparing our previously employed drug screening assay workflow^4^ and a new, refined workflow. **a**, Graphical summary of main steps in drug screening assay according to our previous publication^4^ (“One Pot” Differentiation). **b,** New workflow including an additional step that comprises the differentiation of cells in a separate dish before being seeded into the final assay plate (“Separate Pot” Differentiation). This ensures less variability in the cell count per well and thus higher reproducibility with lower variance in the results.

**Extended Data Fig. 9:**
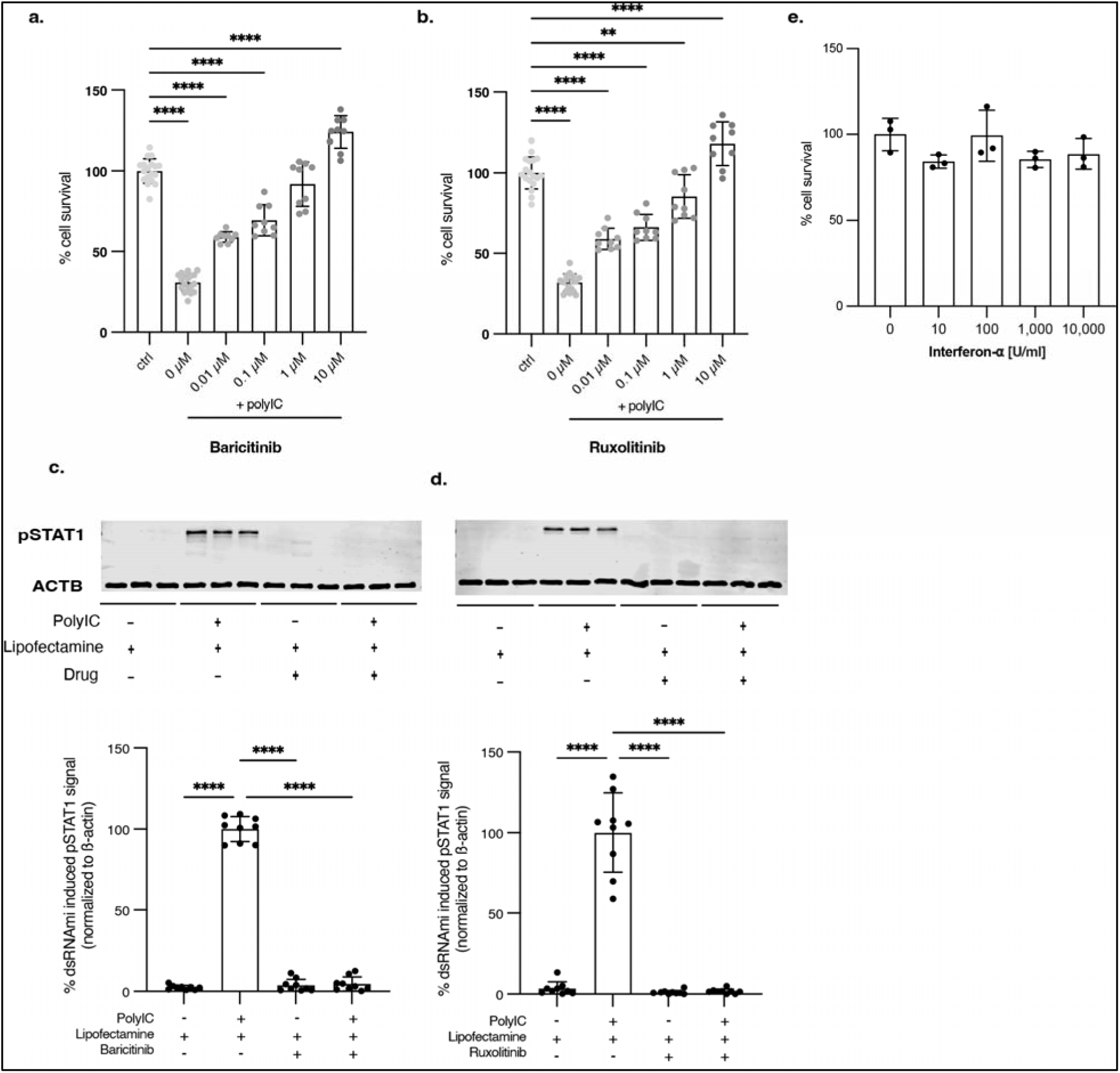
Further validation of baricitinib and ruxolitinib in refined workflow as well as validation of IFNAR2 as a drug target. **a-b**, Quantification of cell survival of ReN VM cell-derived neurons pre-treated with baricitinib (a) or ruxolitinib (b) at different concentrations (baricitinib: 0 µM *P* < 0.0001, n = 18; 0.01 µM *P* < 0.0001, n = 9; 0.1 µM *P* < 0.0001, n = 9; 1 µM *P* = 0.0872, n = 9; 10 µM *P* < 0.0001, n = 9; ruxolitinib: 0 µM *P* < 0.0001, n = 18; 0.01 µM *P* < 0.0001, n = 9; 0.1 µM *P* < 0.0001, n = 9; 1 µM *P* = 0.0016, n = 9; 10 µM *P* < 0.0001, n = 9) and afterwards transfected with poly(I:C) or lipofectamine as a vehicle control (ctrl, n = 18). Baricitinib: EC50 = 35.7 nM; ruxolitinib: EC50 = 47.7 nM. **c-d,** Image (top) and quantification (bottom) of Western blot of pSTAT1^Y701^ in ReN VM cell-derived neurons 24h after treatment with baricitinib (c) or ruxolitinib (d) and transfection with poly(I:C) normalized to the housekeeping protein beta actin (ACTB). Top: n = 3 per condition. Bottom: n = 9 per condition (all *P* < 0.0001). **e,** Quantification of cell survival of ReN VM cell-derived neurons treated with different doses of interferon-α (n = 3 per condition). The error bars represent the standard deviation.

