## Supplementary material for "TYK2 as a novel therapeutic target in Alzheimer’s Disease with TDP-43 inclusions": SI Guide

**Supplementary Information Guide**

**Supplementary Table 1: RNA-sequencing data analysis looking for upregulated ISGs in Alzheimer’s Disease brains**

Excel table showing the results of a differential gene expression analysis in Alzheimer’s Diseases patients compared to healthy controls (Fig. 1d, Extended Data Fig. 3). RNA-sequencing data derived from ROSMAP and MSBB were used to evaluate which interferon-stimulated genes were differentially expressed in selected brain regions relevant in Alzheimer’s Disease.

**Supplementary Table 2: CRISPR screen data**

Excel table showing the results of a CRISPR screen in ReN VM-derived neural cells using the Brunello library (Fig. 3a,b).
